# Characterisation of FADD interactome reveals novel insights into FADD recruitment and signalling at the Death Inducing Signalling Complex (DISC)

**DOI:** 10.1101/2021.03.25.436271

**Authors:** Joanna L Fox, Laura S Dickens, Rebekah Jukes-Jones, Gareth J Miles, Claudia Langlais, Kelvin Cain, Marion MacFarlane

## Abstract

Fas-associated death domain protein (FADD) plays a vital role in the extrinsic apoptotic pathway, where it forms an essential component of the death-inducing signaling complex (DISC). However, the precise early molecular events that facilitate recruitment of FADD to the DISC remain poorly defined. Using affinity purification and mass spectrometry we investigated the FADD interactome in untreated cells and following death receptor stimulation to identify novel FADD-interacting proteins. As expected, in death receptor-stimulated samples our analysis identified key components of the DISC such as Caspase-8. In addition, we identified novel binding partners including Transferrin Receptor 1 (TfR1) and Myosin Light Chain Kinase 2 (MYLK2) that are able to modulate FADD recruitment to the DISC and consequently downstream apoptotic signaling. TfR1 is pre-associated with FADD and recruited into the DISC; moreover, our data reveal that TfR1 is also pre-associated with the death receptors, TRAIL-R1 and TRAIL-R2, thereby functioning as a key regulator of DISC formation. In the case of MYLK2, specific binding of FADD to MYLK2 in non-apoptotic cells sequesters FADD from other DISC components ensuring aberrant apoptosis is not initiated. Furthermore, MYLK2 enzymatic activity is required to for it to translocate, in complex with FADD, to sites of DISC-mediated death receptor oligimerization. Taken together, our study highlights the important role that additional novel FADD binding partners play in the regulation of death receptor-mediated apoptotic cell death, in part by modulating FADD recruitment to the DISC.

## Introduction

Commitment of a cell to programmed cell death or apoptosis is a tightly controlled process and has been studied in depth because of the possible consequences its de-regulation can have on human health. There are two main apoptotic pathways; one triggered by binding of ligands to death receptors at the cell surface (extrinsic apoptosis) and the other initiated by signals internal to the cell such as DNA damage (mitochondrial or intrinsic apoptosis).

Extrinsic apoptosis is mediated through death receptors, which are members of the tumour necrosis factor receptor superfamily characterized by a cytoplasmic region known as the “death domain” (DD). The most extensively studied members of this family of receptors are TNF-receptor 1 (TNF-R1), CD95 (Fas/Apo1), TRAIL-R1 and TRAIL-R2. Upon binding of their respective cognate ligands, death receptors trimerise (Kischkel et al., 1995; Siegel et al., 2004) and the adaptor protein FADD is recruited to the DD of the receptor (Boldin et al., 1996; Chinnaiyan et al., 1995). Caspase-8/10 is then able to bind to the Death Effector Domain (DED) of FADD facilitating formation of the Death Initiating Signalling Complex (DISC) at the cell membrane (Dickens et al., 2012; Schleich et al., 2012). Overexpression of the DD of FADD alone (Newton et al., 1998; Newton et al., 2001), mutation of key residues within the CD95-DD (Huang et al., 1996; Martin et al., 1999) mutation of FADD DED domain (Hughes et al., 2016) blocks DISC formation, highlighting the essential role FADD plays as an adaptor protein facilitating recruitment of pro-caspase-8/10 and DISC formation. Proteomic (LC-MS/MS) analysis of the stoichiometry of the native CD95/TRAIL DISC isolated from cells revealed that FADD was present at sub-stoichiometric levels, compared to both death receptors and Caspase-8 (Dickens et al., 2012; Schleich et al., 2012). This, taken together with the observed ‘death effector filaments’ visualised by confocal microscopy in cells expressing GFP-labelled Caspase-8 tandem DEDs (tDEDs) (Dickens et al., 2012; Siegel et al., 1998) led to the proposal of the helical chain model. Whereby, once the complex is nucleated by FADD recruitment to the receptor, multiple procaspase-8 molecules bind sequentially primarily via a FL:Pocket interaction between tDEDs (Dickens et al., 2012). Subsequent structural analysis of the Caspase-8 tDED filament by Cryo-electron microscopy (Fu et al., 2016), revealed that three Caspase-8 tDED strands interact to form a triple helix structure. Once this filament structure has formed via the tDEDs of Caspase-8, the catalytic domains of pro-caspase-8/10 are able to dimerise in an anti-parallel orientation (Fox et al., 2021; Watt et al., 1999) and proteolytically cleave key residues resulting in caspase activation (Blanchard et al., 1999; Pop et al., 2007; Wang et al., 2010b). These cleavage events result in the initiation of the apoptotic cascade that drives commitment to cell death. The order of early events upon CD95 receptor activation resulting in DISC formation and commitment to apoptosis has previously been established (Algeciras-Schimnich et al., 2002) revealing that FADD and Caspase-8 recruitment to the receptor occur within a few minutes of ligand binding. However, receptor clustering, and in Type 1 cells subsequent receptor internalization, occur as distinct steps in the initiation process. Interestingly, both the recruitment of key DISC components and CD95 receptor internalization were found to be dependent on the actin cytoskeleton, inhibition of which blocked both DISC formation and subsequent commitment of the cell to apoptotic cell death (Algeciras-Schimnich et al., 2002).

However, the key to all these processes is the adaptor protein FADD which, although initially described in death receptor signalling, has subsequently been found to be involved in numerous other signalling settings including embryonic development, cell survival, proliferation, division, tumor progression, TLR-signaling, inflammation, innate immunity, necrosis, and autophagy (Tourneur and Chiocchia, 2010). FADD has two distinct protein binding domains, the DD at the N-terminus and the DED at the C-terminus that enables it to function as an adaptor protein. The DD of FADD interacts with other DD containing proteins such as the death receptors TRAIL-R1/R2 and FAS, whereas the DED facilitates interaction with DED containing proteins such as Caspase-8, Caspase-10 and c-FLIP. Recruitment of FADD into these various signalling complexes, particularly the DISC acts as a nucleating event, after which further proteins are recruited to form the active complex. Interestingly, both the subcellular localisation (Gomez-Angelats and Cidlowski, 2003) and phosphorylation status (Alappat et al., 2005) of FADD appear to be key in determining the complex that forms and consequently its function in the cell. There have been numerous studies to determine, at a structural level the interaction between FADD DD and CD95 which have provided insights into how the DISC assembles (Esposito et al., 2010; Scott et al., 2009; Wang et al., 2010a). However, despite these substantial advancements in understanding the process of death receptor initiated apoptosis the early molecular events that facilitate recruitment of FADD to the DISC are not fully understood.

In this study we have purified FADD-containing multi-protein complexes pre- and post-death receptor stimulation and used label-free quantitative LC-MS/MS to identify previously unknown FADD-interacting proteins. Using this approach, we identified Transferrin Receptor 1 and Myosin Light Chain Kinase 2 (MYLK2) that are able to modulate FADD recruitment to the DISC and as a result downstream apoptotic signaling. Furthermore, our data reveal how FADD is regulated in non-apoptotic cells to ensure aberrant apoptosis does not occur, whereas once apoptosis is initiated it is able to translocate to sites of death receptor trimerization to form the DISC. Thus, we now provide new insights into the early activation events that facilitate DISC formation and activation.

## Results

### FADD interacts with multiple protein complexes both in untreated and CD95 treated cells

In this study, we aimed to identify binding partners of FADD both before and after initiation of DISC formation triggered by anti-CD95. FADD-/- Jurkat cells were transfected with vectors expressing a FADD-TAP C-terminal affinity construct (Fig 1A) and single clone cell lines established. Two of the clones characterized had FADD expression levels similar to wild-type (WT) Jurkat A3 cells (Fig 1B). Importantly, re-expression of FADD-TAP had no detectable effect on the expression of other key apoptotic proteins, including Caspase-8, −3 and −9, the DISC regulator FLIPL/S, or mitochondrial apoptotic regulatory proteins BCL-2, BID or BAK (Fig 1B).

**Figure 1.**
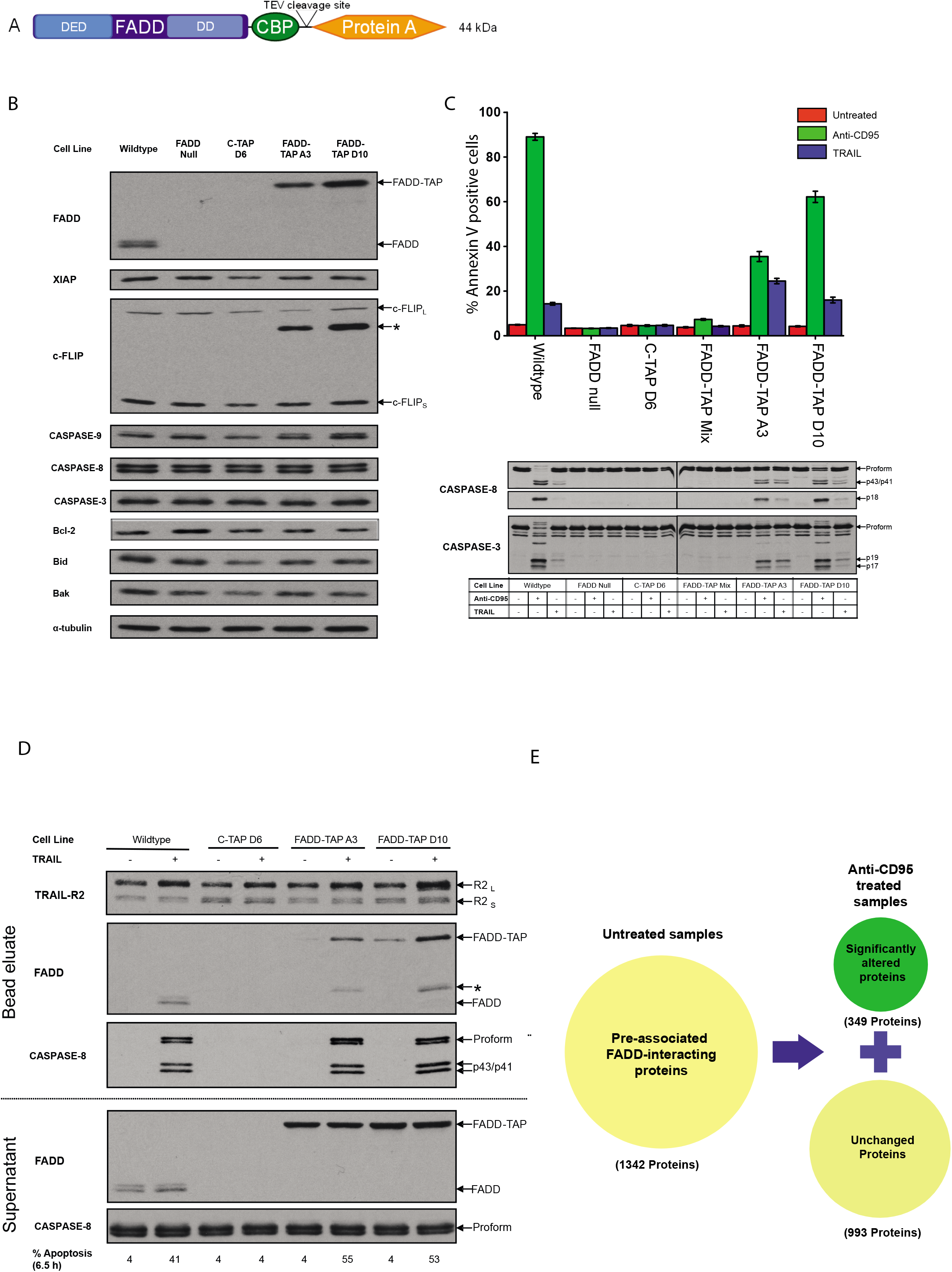
FADD-TAP re-expression in FADD-null cells restores sensitivity to anti-CD95 and TRAIL-mediated apoptotic cell death. A) Schematic representation of FADD-TAP construct. B) Western blot analysis of FADD-TAP expression in Jurkat FADD^-/-^ cells, along with other key apoptotic proteins. C) Apoptotic response to anti-CD95 (green bars) or TRAIL (blue bars) treatment on wild-type, FADD-null, empty vector (C-TAP D6) or FADD-TAP transfected clones (mixed, A3 or D10) compared to untreated (red bars) cells, as determined by Annexin V/Draq7 staining and corresponding Caspase-8 and −3 cleavage visualized by western blotting. D) Western blot analysis of TRAIL DISC, stimulated with bTRAIL and isolated by affinity purification. DISC-containing eluates and cleared lysate supernatants (1% of total input) were probed for the known components. Prior to lysis 1×10^6^ cells were kept at 37°C for 6.5 h for assessment of apoptosis by Annexin V/Draq7 staining. The values show percentage apoptosis (Annexin V positive cells). E) Schematic representation of the FADD-interacting protein subgroups identified by LC-MS/MS.

Treatment of the FADD -/- parental cells and C-TAP D6 empty vector single clone control cell line, with anti-CD95 or TRAIL did not induce apoptotic cell death as determined by Annexin V/Draq7 staining and there was no detectable cleavage of Caspase-8 or −3 (Fig 1C). The mixed FADD-TAP (unsorted) population exhibited negligible levels of cell death and no detectable Caspase-8/-3 cleavage. However, in the single clone cell lines (A3 and D10), FADD re-expression resulted in apoptotic cell death in response to both ant-CD95 and TRAIL treatment, although the levels of cell death were lower than in WT cells. Furthermore, in WT, A3 and D10 cells, higher levels of cell death were induced by anti-CD95 as compared to TRAIL, with cleavage of both pro-caspase-8 and −3 detected in response to both ligands (Fig 1C). Also, in the re-expressed FADD-TAP cells (A3 and D10), FADD was recruited to the DISC at similar levels to that seen in WT cells, resulting in Caspase-8 activation as determined by cleavage of procaspase-8 to its p43/41 form (Fig. 1D).

Having established that expression of FADD-TAP in FADD-null cells restored anti-CD95 -mediated cell death, we then used affinity purification to identify FADD-interacting proteins pre- and post-DISC formation. Empty vector (D6) or FADD-TAP expressing (D10) cells were incubated on ice for 1h +/- anti-CD95 prior to incubation at 37°C for 20 min to trigger DISC formation, but not allow the cells to progress to apoptotic cell death. FADD and any binding partners were then purified by affinity purification via the Protein-A tag on FADD using IgG agarose, and co-purified proteins identified using label-free quantitative mass spectrometry. Proteomic analysis was performed on FADD-TAP and empty vector expressing cells both pre- and post-CD95-induced DISC formation. FADD is known to have both apoptotic and non-apoptotic roles within the cell, therefore by isolating all FADD from the cells, and not specifically FADD within the DISC, we identified three groups of proteins interacting with FADD. Those pre-associated with FADD, anti-CD95-induced interacting proteins that either associate or dissociated from FADD following anti-CD95 treatment, and pre-existing FADD-interacting proteins unaffected by triggering of DISC formation (Fig. 1E). To remove non-specific interacting proteins from the dataset, any proteins that were present in the empty vector control samples were considered to be non-specific contaminants that were deleted from the analysis. We subsequently identified 1342 proteins that were interacting with FADD-TAP, 79% of which were not significantly altered following anti-CD95-induced DISC formation (Fig. 2A). Of the 21% of proteins identified as significantly changed, nearly 100% were significantly increased, with only 1 protein significantly decreased (Fig. 2B). Those proteins, which increased upon anti-CD95 treatment, were further classified by function using GO annotation; consequently, some proteins are assigned to multiple ontologies (Fig. 2C). Interestingly, proteins from a whole range of different processes were altered after death receptor stimulation including those involved in cell death (Fig. 2C) with no one cellular function standing out as being highly enriched. We were able to identify some known FADD-interacting proteins in our data set. Caspase-8 was only detected following anti-CD95 treatment, whereas Casein kinase 2 (CSK22/23) was identified in untreated samples and the amounts detected decreased following treatment (Fig. 2D). Interestingly, we also observed that the amount of FADD detected was decreased following anti-CD95 treatment. In addition, we detected numerous components of the proteasome in our data (Fig. 2E), which correlates with previous reports of proteasome-induced degradation of FADD following DISC formation to terminate signaling (Lee et al., 2012).

**Figure 2.**
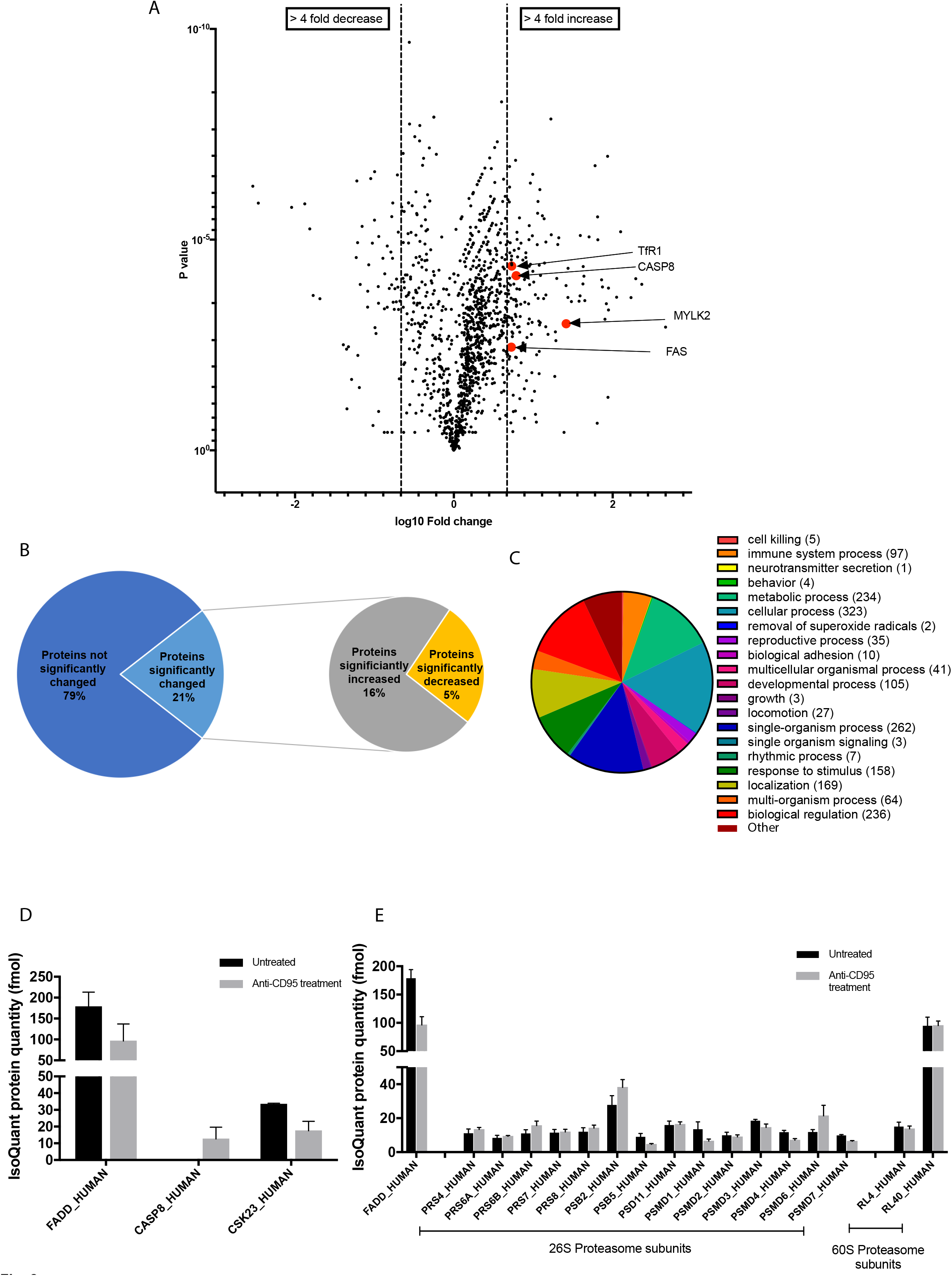
FADD interacts with multiple protein complexes, both in untreated and anti-CD95 treated cells. A) Volcano plot of FADD-TAP interacting proteins identified by label-free LC-MS/MS; >4 fold-change in amount of proteins following anti-CD95 treatment was determined as significantly altered. B) Graphical representation of percentage of proteins identified by mass spectrometry that were significantly changed following anti-CD95 treatment. This was further divided into those proteins that were up or down -regulated. C) Pie chart of gene ontology pathway analysis of proteins identified by mass spectrometry. D & E) Label-free quantitation of amount of each protein detected +/- anti-CD95 treatment (n=3 biological repeats, Mean +/- SEM).

### FADD and TRAIL-R1/R2 associate with Transferrin Receptor 1 both prior to and during DISC formation

Transferrin receptor 1 (TfR1) was consistently detected both pre-associated with FADD as well as with the CD95 DISC (Suppl Fig. 1). Furthermore, the interaction of TfR1 was enhanced following DISC stimulation. Interestingly, in our previous label-free LC-MS/MS analysis of the stoichiometry of the TRAIL DISC (Dickens et al., 2012)(Dickens et al, 2012), we had also detected TfR1 as a novel component of the TRAIL DISC (Fig. 3A). Thus, to further investigate the presence of TfR1 at the DISC we isolated and analyzed the composition of the DISC using Biotin-TRAIL. In multiple cell lines expressing both TRAIL-R1 and TRAIL-R2, we were able to detect TfR1 both pre-associated with the receptors when biotynlated TRAIL was added to lysates from untreated cells (TRAIL postlysis), and also in the DISC isolated from TRAIL-treated cells. (Fig. 3B). Further analysis of TRAIL-treated BJAB cell lysates by sucrose density gradient, which allows complexes to be separated by buoyant density prior to affinity purification and analysis by western blotting, revealed that TfR1 is found in a complex with TRAIL receptors. Analysis of the high molecular weight DISC confirmed the presence of TfR1 in the DISC, in this case triggered by treatment of cells with TRAIL (Fig. 3C). As TfR1 is pre-associated with the death receptors, we investigated whether TfR1 has a role in trafficking the receptors to the cell surface (Fig. 3D). To further investigate the role of the TfR1-TRAIL-R1/2 complexes we knocked down TfR1 using siRNA. Knock down of TfR1 had no effect on surface expression of TRAIL-R1 but did increase the surface expression of TRAIL-R2, compared to the transfection and non-targeting siRNA controls. Importantly, knockdown of TfR1 reduced the amount of Caspase-8 activation following TRAIL treatment, as evidenced by the decrease in Caspase-8 cleavage observed (Fig. 3E). This consequently resulted in less mitochondrial outer membrane permeabilisation (Fig. 3F) and decreased caspase-7 cleavage (Fig. 3E), suggesting that in these cells, Caspase-8 is unable to cleave and activate BID resulting in less apoptosis *via* the intrinsic apoptosis feedback loop. This reduction in TRAIL-mediated Caspase-8 activation, was due to a decrease in DISC formation as shown by decreased levels of TRAIL-R1/-R2, FADD and Caspase-8 within the DISC (Fig. 3G). Taken together these data suggest that TfR1 is pre-associated with both FADD and death receptors and plays a role in formation of a functional DISC. Furthermore, TfR1 pre-association with critical components and its subsequent recruitment to the DISC is a general mechanism for death receptors.

**Figure 3.**
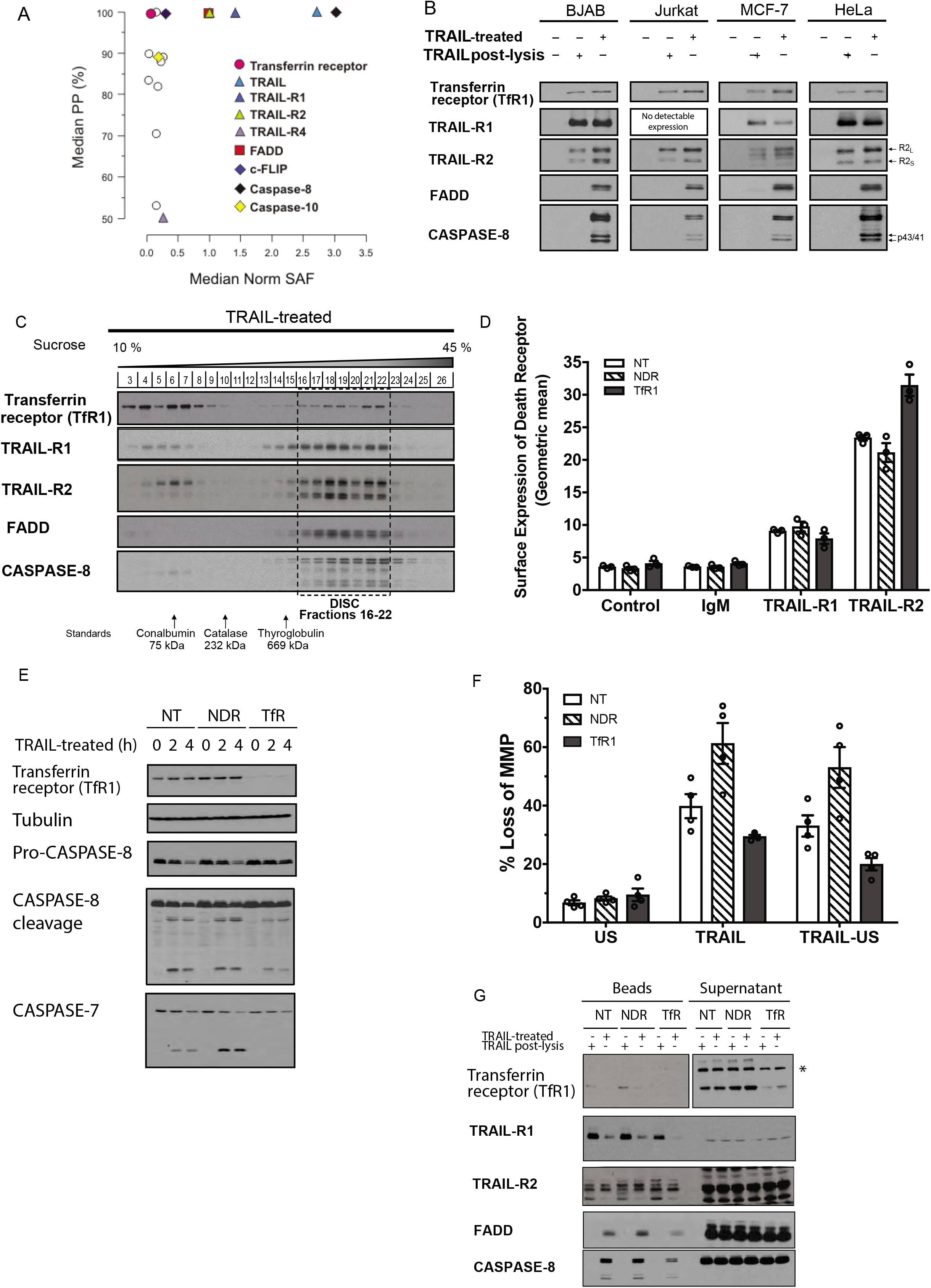
FADD and TRAIL-R1/R2 associate with Transferrin Receptor 1, both prior to and during DISC formation across multiple cell lines. A) Graphical analysis of TRAIL DISC mass spectrometry (Dickens et al., 2012). Proteins identified by LC-MS/MS analysis of TRAIL DISC purified from BJAB cells were plotted according to mean protein probability (PP) and median spectral abundance factor (SAF) relative to TRAIL-R2 (n=3 biological repeats). B) Western blot analysis of Transferrin Receptor 1 (TfR1) in TRAIL-DISC isolated from BJAB, Jurkat, MCF-7 and HeLa cells. TRAIL added post-lysis enabled isolation of unstimulated TRAIL-R1/R2. C) Western blot analysis of bTRAIL stimulated DISC formation. Protein complexes from BJAB cell lysates treated +/- 500ng/ml bTRAIL were first separated by buoyant density on 10-45% sucrose density gradient, then TRAIL-DISC isolated by affinity purification. The dashed boxed area indicates fractions that contain the DISC. Molecular weight standards were run on a separate gradient and the positions shown are representative of three independent experiments. Note, in unstimulated BJAB cell lysates, TfR1 elutes at it’s native molecular weight of ~80kDa (see Suppl Fig. 2). D) Determination of surface expression of TRAIL-R1 and TRAIL-2 following RNAi knockdown of TfR1, or control samples non-targeting RNAi (NT), and RNAi against a non-Death Receptor protein (NDR) in MCF-7 cells. E) Western Blot analysis of effect RNAi knockdown of Transferrin Receptor 1 (TfR1), or control non-targeting RNAi (NT), and RNAi against a non-Death Receptor protein (NDR) on Caspase-8 and −7 cleavage in MCF-7 cells after treatment with TRAIL for 0, 2 or 4 h. F) Effect of RNAi knockdown of TfR1, or control non-targeting RNAi (NT), and RNAi against a non-Death Receptor protein (NDR) on induction of mitochondrial membrane permeabilisation (loss of MMP) in MCF-7 cells following treatment with TRAIL for 2 h. G) Western blot analysis of TRAIL-DISC isolated from MCF-7 cells, following siRNA with a non-targeting RNAi (NT), RNAi against a non-Death Receptor protein (NDR) or transferrin receptor (TfR1).

### FADD associates with cytoskeletal protein MYLK2 both prior to and within the DISC

We detected numerous proteins involved in the cytoskeleton both pre-associated with FADD and present following DISC induction (graph shows mean protein amounts detected in 3 independent biological repeats; Fig. 4A). Of these proteins, we observed Myosin Light Chain Kinase 2 (MYLK2) to be consistently pre-associated with FADD and up-regulated following anti-CD95 treatment (Fig. 2A & 4B). Western blot analyses confirmed MYLK2 bound to purified FADD-TAP (FADD-TAP D10) but not to the beads from the empty vector (C-TAP D6) cells, validating the mass spectrometry data (Fig. 4C). To confirm the interaction between MYLK2 and FADD was not an artifact of the overexpression of the FADD-TAP construct, we immunoprecipitated native FADD from Jurkat (Suppl Fig. 3A) and BJAB cells (Suppl Fig. 3B) cells and probed for the presence of MYLK2. This confirmed the pre-association of MYLK2 with FADD in untreated cells. We then utilized two complementary approaches to determine whether MYLK2 was recruited with FADD to the DISC as suggested by the observed enrichment in the mass spectrometry data (Fig. 4A and 4B). Firstly, FADD-TAP cells were treated with anti-CD95 and the complexes that formed separated by size exclusion chromatography. We were clearly able to identify the DISC components (CD95/Caspase-8/FADD) in a discrete number of fractions. MYLK2 is also present in these fractions (Fig 4D). It is also interesting to note that not all the FADD protein is found within the DISC (fractions 5-13), indicating only a subset of FADD is recruited to the complex when a cell is treated with anti-CD95. We further confirmed recruitment to the native CD95 DISC by immuneprecipitating the complex using biotin-anti-CD95, where along with the expected components FADD, Caspase-8 and CD95, MYLK2 was also present in the DISC (Fig 4E).

**Figure 4.**
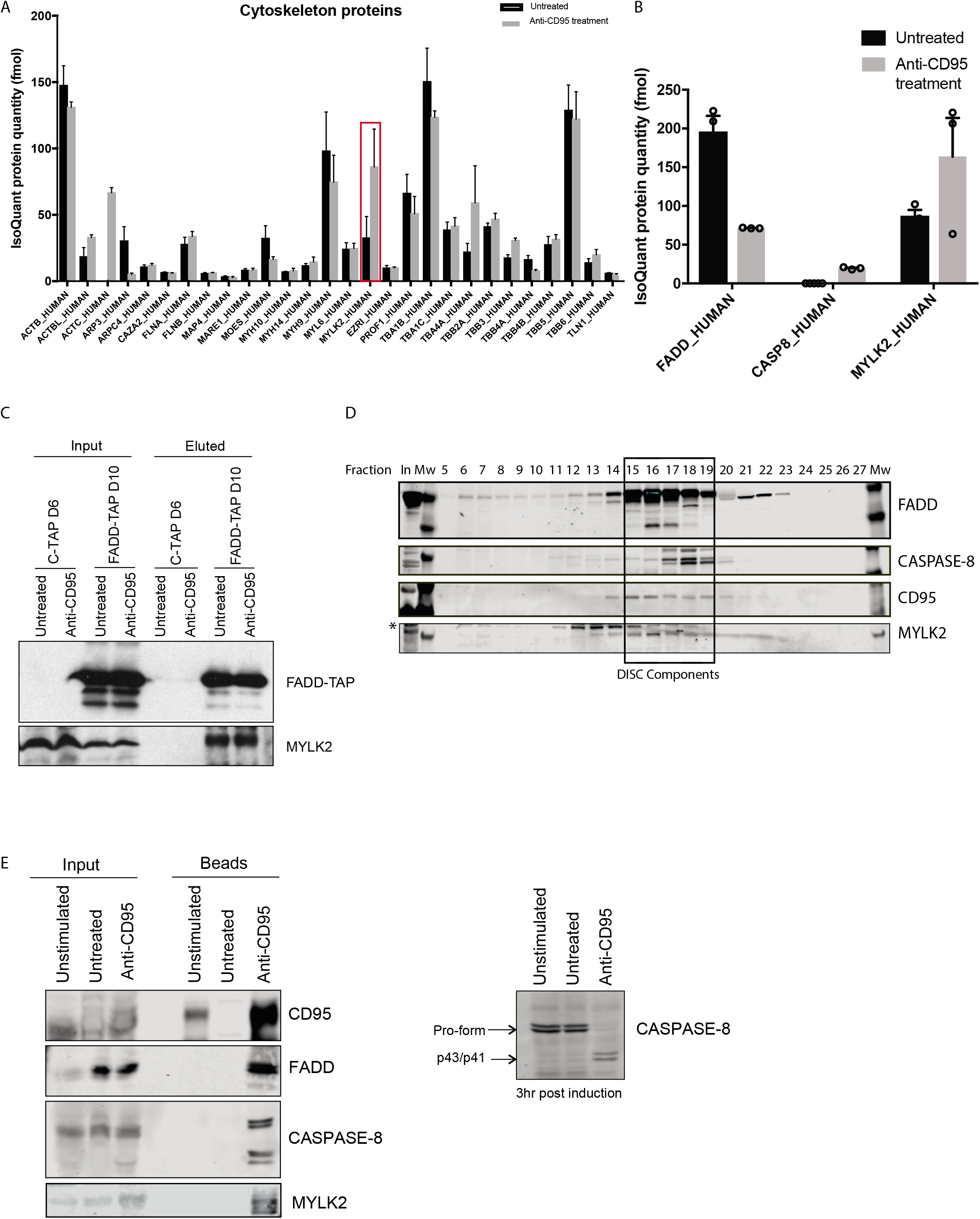
FADD interacts with cytoskeletal protein MYLK2 both prior to and during CD95 DISC formation. A) Label-free LC-MS/MS quantitation of cytoskeletal-related proteins detected +/- anti-CD95 treatment (n=3 biological repeats, Mean +/- SEM). B) Label-free LC-MS/MS quantitation of FADD and known/novel FADD-interacting proteins detected +/- anti-CD95 treatment (n=3 biological repeats, Mean+/- SEM). C) Western blot analysis of affinity purified TAP-tag complexes from empty vector (C-TAP D6) and FADD-TAP cells (FADD-TAP D10). D) Superdex 200 gel filtration fractions of whole cell lysates from FADD-TAP cells following anti-CD95 DISC formation followed by western blotting for FADD, Caspase-8, CD95 and MYLK2. *denotes non-specific bands; Mw = Molecular weight markers; In = Input whole cell lysate loaded onto column. E) Western blot analysis of CD95 DISC isolated from BJAB cells. Prior to lysis, 1×10^6^ cells were kept at 37°C for a further 3 h for assessment of anti-CD95-induced apoptosis by Caspase-8 cleavage.

### RNAi knockdown of MYLK2 reduces sensitivity to CD95L–induced apoptosis

We used siRNAi to knockdown MYLK2 levels to determine if this affected the apoptotic function of FADD. We achieved approximately 50% knockdown in BJAB cells 24 h post-transfection (Fig. 5A). However, this level of knockdown was sufficient to reduce CD95L-induced apoptosis after 4 h of treatment, as determined by Annexin V/Draq7 staining (Fig. 5B). It is known in Type I cells such as BJABs that following receptor-ligand activation the receptor is internalized and the total level of surface expression of the receptor is reduced (Algeciras-Schimnich et al., 2002). This can be measured by labeling any exposed receptor on the surface of cells with a fluorescently-labeled antibody and measuring the read-out by flow cytometry. As has been previously reported, when control cells both with and without nucleofection were treated with CD95L for 30 min the level of surface-exposed CD95 receptor was reduced. However, when cells, which had MYLK2 levels reduced *via* RNAi, were similarly treated with CD95L no change in the surface expression of CD95 receptor was detected (Suppl Fig. 4), suggesting that a functional DISC is not formed when MYLK2 levels are reduced. This, however, could not be confirmed at the level of the DISC, as protein knockdown by RNAi was only feasible in a small number of cells.

**Figure 5.**
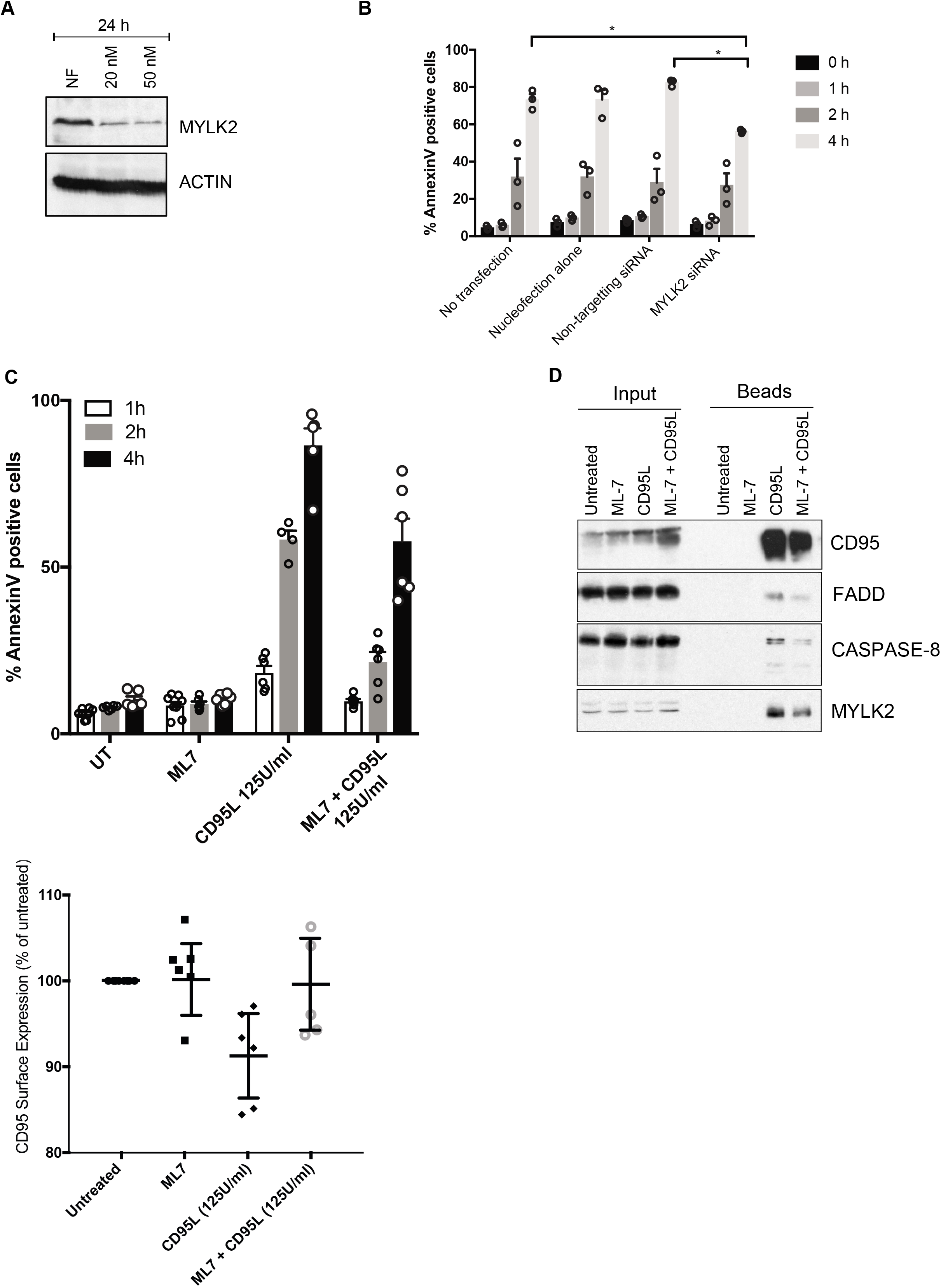
RNAi knockdown or Inhibition of MYLK2 reduces sensitivity to CD95L-induced apoptosis and prevents FADD recruitment to the DISC. A) Western blot analysis of RNAi knockdown of MYLK2 in BJAB cells. NF = nucleofection control; MYLK2 siRNA (20 nM, 50 nM). B) Time course of apoptotic response to CD95L treatment following 24h RNAi knockdown of MYLK2 in BJAB cells as determined by Annexin V/Draq7 staining (n=3 biological repeats, Mean +/- SEM). C) Time course of apoptotic response to CD95L treatment +/- pre-treatment with the MYLK2 inhibitor, ML7, in BJAB cells as determined by Annexin V/Draq7 staining. D) Western blot analysis of CD95 DISC isolated from BJAB cells +/- ML7 pre-treatment. E) Determination by FACS analysis of surface expression of CD95 following 30 min CD95L treatment +/- ML7 pre-treatment. Surface expression expressed as a percentage compared to the untreated control (duplicate samples from 3 biological repeats, Mean +/- SEM).

### Inhibition of MYLK2 decreases apoptosis in response to CD95L and prevents FADD recruitment to the DISC

To determine whether MYLK2 had a scaffold role or whether the kinase activity of MYLK2 was involved in the anti-apoptotic effect we observed, we treated cells with the MYLK2 inhibitor, ML7. Treatment of cells with this compound for 2 h did not cause any significant cell death in the cells as determined by Annexin V/Draq7 staining (Fig. 5C). However, when cells were pre-treated with ML7 for 30 min prior to treatment with CD95L, there was a significant decrease in the cell death observed (Fig. 5C). As we had observed MYLK2 within the DISC, we investigated the effect of MYLK2 inhibitor on DISC formation. The DISC was not detected in untreated cells or cells treated with ML7 alone. As expected, cells treated with biotin-anti-CD95 alone or ML7 (pre-treated) combined with biotin-anti-CD95 formed a DISC and in both cases contained MYLK2 along with CD95, FADD and Caspase-8. However, we consistently detected less FADD, and consequently reduced Caspase-8 in cells pre-treated with ML7 (Fig. 5D). There was also less CD95 and MYLK2 compared to the untreated samples suggesting that CD95 DISC formation is less efficient in the absence of MYLK2 kinase activity. CD95 is known to internalize following DISC initiation resulting in lower levels of receptor detected at the cell surface, so we measured the level of CD95 on the surface of cells following DISC stimulation by CD95L with and without inhibition of MYLK2. ML7 alone had no affect compared to untreated cells, whereas following CD95L treatment the level of CD95 receptor detectably decreased. Interestingly, in cells pre-treated with ML7 prior to DISC stimulation with CD95L, there was less CD95 receptor internalization (Fig. 5E). Taken together with the reduced levels of FADD and Caspase-8 in the CD95 DISC isolated from ML7 pre-treated cells, the kinase activity of MYLK2 is clearly required for efficient CD95 DISC formation, internalization and initiation of cell death.

## Discussion

The importance of correct DISC assembly is well established (Esposito et al., 2010; Fox et al., 2021; Keller et al., 2009; Scott et al., 2009; Wang et al., 2010a). In this study, we aimed to identify new FADD-interacting proteins involved in both the DISC and regulation of the assembly process. Our study has highlighted the diverse array of functions that FADD is involved with in the cell. The protein hits identified by mass spectrometry are involved in many different cellular processes (Fig. 2B). Validation of this approach and potential insight into the role of FADD in DISC regulation was confirmed by our ability to identify previously characterized pathways for DISC regulation. Importantly, we were able to identify Caspase-8 recruitment only following anti-CD95 treatment and were also able to find Casein kinase 2 reproducibly in untreated samples, a kinase which is reported to phosphorylate FADD in non-apoptotic cells (Vilmont et al., 2015). We identified proteins that are pre-associated with FADD prior to CD95 stimulation, and those proteins that only associate once DISC formation has been triggered. The pre-associated proteins could be further sub-divided into those whose interactions are unchanged, increased or decreased following anti-CD95 treatment. Proteins whose interactions were unchanged as a result of anti-CD95 treatment were the largest group of proteins within the analysis; thus, highlighting an important fact that when DISC formation is triggered this does not involve all cellular FADD being recruited. Rather, that FADD has a high-affinity group of proteins it remains associated with in non-apoptotic cells; this is likely to allow FADD to facilitate its non-apoptotic roles in the cell (Alappat et al., 2005; Werner et al., 2006). The consequence of FADD remaining bound to these other partner proteins could be that it is essentially sequestered and thus unavailable for recruitment to the DISC, providing a possible explanation for why only a small proportion of total FADD is recruited to the DISC.

Another mechanism of DISC regulation is via FADD turn-over within the cell by the Ubiqutin-proteasome pathway (Lee et al., 2012). It was suggested that the magnitude and duration of the apoptotic response to initiation of extrinsic signaling was determined by FADD. Furthermore, once FADD has been recruited to the DISC and apoptosis successfully initiated, the signal needs to be negatively regulated, which is achieved by protein degradation. However, in the context of this study, although total FADD levels have been decreased apoptosis has not been allowed to proceed - thus, any proteins recruited with FADD to the DISC should be largely unaffected by this pathway. It is likely that FADD being targeted for degradation, in this case is not bound into a multi-protein complex.

The first new FADD interacting protein we identified in this study was Transferrin Receptor 1 (TfR1), which was found associated with FADD both before and after anti-CD95 treatment. Further investigation revealed that TfR1 was also pre-associated with the death receptors TRAIL-R1 and TRAIL-R2. TfR1 is a membrane receptor, which is involved in the uptake of iron and upregulation of TfR1 protects cancer cells (Fassl et al., 2003; Habashy et al., 2010) against death receptor mediated cell death. Conversely TfR1 has been implicated in gambogic acid-induced apoptosis, *via* activation of Caspase-8 and the intrinsic cell death pathway, but not through an association with known DISC components (Kasibhatla et al., 2005). To date however, this is the first report of a pre-association between TfR1 and either FADD or TRAIL-R1/TRAIL-R2 prior to initiation of apoptotic signalling. It still remains to be defined whether these interactions are direct or indirect as TfR1 lacks the CRD domain that mediates death receptor interactions (Clancy et al., 2005; Naismith and Sprang, 1998), therefore, any interaction between these proteins must occur *via* their cytoplasmic domains. One possibility therefore, is that binding of both death receptors and FADD to TfR1 is a way of sequestering DISC components prior to any apoptotic signal to prevent spontaneous interactions between the components. Interestingly, a recent study correlating TfR1 expression and survival in astrocytic brain tumours revealed that high expression of TfR1 was associated with poor overall survival (Rosager et al., 2017). Although this study did not explore levels of apoptosis in the patient samples, it would be interesting to determine if the high TfR1 expression affected the apoptotic threshold of the cells, by preventing the initiation of apoptosis by limiting DISC formation.

We identified Myosin light chain kinase 2 (MYLK2) as another new pre-associated FADD binding partner prior to apoptotic induction, which after death receptor activation translocated with FADD into the DISC. This skeletal or short isoform of the kinase encodes the core kinase domain, plus a regulatory domain that includes auto-inhibitory and Ca2+/Calmodulin-binding sequences. MYLK2 also contains an actin binding sequence that tethers it to the cytoskeleton. With increasing evidence that cytoskeletal proteins may participate in the regulation of cell survival and apoptosis, the identification of MYLK2 as a strong FADD binding protein is intriguing. Activation of the long isoform of MYLK2 has been reported to be an early step in signalling via the TNF complex and triggers the recruitment of TNFR1 binding proteins including TRADD, TRAF2, the adaptor protein RIP1, plus other yet unknown proteins (Jin et al., 2001). However, to date this is the first report of the involvement of the short isoform of MYLK2 in apoptosis signalling. Most studies focus on the role of MYLKs in phosphorylating myosin light chain to regulate actomyosin contractility (Stull et al., 2011). However, there are some reports characterising a scaffolding role for MYLK not requiring its catalytic activity, and an involvement in regulation of the cytoskeleton by recruitment of specific kinases or cortactin to specific cellular compartments (Dudek et al., 2010; Usatyuk et al., 2012). Our study shows that there is a threshold amount of active of MYLK2 required for maximal FADD recruitment to the DISC, as total cellular apoptosis was reduced either when protein levels were reduced by RNAi or the kinase activity inhibited by the MYLK2 inhibitor, ML-7, suggesting a role for MYLK2 in the translocation of FADD to the receptor.

Calcium ions (Ca^2+^) are important second messengers in cell signaling, able to initiate various cellular responses depending on temporal and spatial parameters. Ca^2+^ release has been observed following CD95 ligand engagement with the receptor, although there are contradictory reports about the role this plays in apoptosis signaling. In some settings, the Ca^2+^ signal was found to be necessary to propagate the FAS induced apoptotic signal (Wozniak et al., 2006). Whereas, other studies report the observed Ca^2+^ release to be anti-apoptotic, by recruiting PKC to the death receptor resulting in inhibition of the apoptotic signal (Khadra et al., 2011). However, all the reports indicate that Ca^2+^ release plays a role early in CD95 signaling. MYLK2 is a Ca^2+^/CaM dependent kinase, therefore, the CD95 induced Ca^2+^ release could contribute to the activation of MYLK2 which is required to facilitate FADD recruitment to the receptor and initiation of DISC formation, although further studies would be required to establish this.

Interestingly mutational analysis in colorectal cancer revealed MYLK2 as one of the eight kinases found to have mutations in the kinase domain (Parsons et al., 2005). Furthermore, low expression in breast cancer correlated with poor prognosis and decreased survival (Kim and Helfman, 2016). In the context of this study, the decreased level of MYLK may impact on the ability of FADD to be recruited to the DISC consequently increasing the apoptotic threshold of the cell and providing the cancer with a survival advantage.

Taken together, our study provides new insight into the diverse roles FADD plays within the cell, as well as determining novel mechanisms of regulation that are essential for the tightly controlled process of recruitment of FADD to the DISC. We now hypothesize that, in non-apoptotic cells FADD is sequestered away from death receptors *via* binding to non-apoptotic proteins such as TfR1 or MYLK2. Once a death ligand binds to its cognate receptor and initiates extrinsic apoptotic signaling there is concurrent influx of Ca^2+^, which could trigger activation of MYLK2 located in proximity to the activated death receptors. Once active, MYLK2 in complex with FADD would be able to translocate *via* the actin cytoskeleton to death receptors at the cell surface and bind into the DISC triggering Caspase-8 recruitment and initiation of apoptosis. Our study therefore highlights the important role the cytoskeleton plays in regulation of apoptotic cell death and the potential impact that reduced FADD recruitment to the DISC has on apoptotic cell death.

## Materials and Methods

### Cell Culture

Four types of Jurkat T cells were cultured. Parental clone A3 (wild-type) and FADD-null Jurkat cells (Juo et al., 1999) were a kind gift from Dr J Blenis (Harvard Medical School, Boston, USA). FADD-TAP and TAP-EASY empty vector Jurkat cells were produced by transfection of FADD-null Jurkat cells with FADD-TAP vectors. FADD-TAP Jurkat cells were selected in 400 μg/ml of the antibiotic G418 (Gibco, Invitrogen). The BJAB cell line was kindly provided by Dr A Thorburn (University of Colorado Health Sciences Center, Aurora, USA (Thomas et al, 2004b)). All cell lines were cultured in RPMI media, supplemented with 10 % v/v FCS and 1 % v/v Glutamax. Cells were routinely cultured at 37 °C with 5 % CO_2_ in a humidified atmosphere.

### Chemicals and Drug Treatments

All chemicals were of the highest quality and were purchased from Sigma (Gillingham, UK) unless otherwise stated and used at the final concentrations indicated. Treatments with ML-7 (20 μM), anti-CD95 (CH11; 500ng/ml), and CD95 Ligand (CD95L; 125U/ml) were carried out for specified times.

### Plasmid Construction

FADD was obtained via PCR amplification of pMal FADD-MBP (Hughes, Mol Cell 2009) and was cloned in-frame into the EcoR1 and Xho1 sites of pcDNA3-C-TAP tag easy (a kind gift from Dr. Tencho Tenev and Prof. Pascal Meier, Institute of Cancer Research, London, UK). The DNA sequence of the vectors was verified by sequencing analysis.

### Western Blotting

Western blot analysis was carried out as previously described (Hughes et al., 2015) Briefly, cells were harvested, washed with PBS, and lysed at 4°C in DISC lysis buffer (30 mM TRIS-HCl (pH 7.5), 150 mM NaCl, 10% glycerol, 1% Triton X-100), containing Complete protease inhibitors. Protein quantitation was carried out using Bradford protein assay (Perbio Science UK Ltd). Samples were run on SDS-PAGE gels and transferred to PVDF membrane. Antibodies used are listed in Table 2. Secondary antibodies were horseradish peroxidase-conjugated goat anti-mouse or goat anti-rabbit (used at dilution of 1:2,000; Dako UK, Ltd.). Reactive proteins were visualized by chemiluminescence with ECL plus (Amersham plc).

### Purification of FADD-TAP and associated proteins

Cells were harvested, washed in PBS, lysed in DISC lysis buffer and centrifuged to remove the insoluble fraction. IgG Sepharose beads were washed with PBS and then with 0.2 M triethanolamine. IgG was cross-linked to the beads with 20 mM dimethyl pimelimidate dihydrochloride for 30 min and the reaction stopped by adding 50 mM Tris-HCl pH 7.5 for 15 min at room temperature. Beads were washed and re-suspended in DISC lysis buffer before addition to the cleared lysate. Lysate and beads were incubated for 1 h at 4°C mixing end-over-end. Beads were removed, washed three times in DISC lysis buffer and proteins eluted from the beads in x2 sample buffer.

### Label-free quantitative mass-spectrometry

FADD and binding partners were isolated as described above. The isolated proteins were separated by SDS-PAGE, using Bolt™ gels. SDS-PAGE gels were sliced into 24 slices, de-stained, dehydrated and digested with trypsin (Promega) overnight at 30°C. The samples from every two adjacent gel slices were combined, to give a total of 12 samples from each lane. Tryptic peptides were extracted with trifluoroacetic acid (TFA; 0.2%), dried in a speedvac then solubilized in 20μl injection solvent (97% FA (5%) 3% MeCN). For Top 3 label-free analysis, the injection solvent was spiked with 20 fmol/μl yeast alcohol dehydrogenase (ADH) and bovine serum albumin (BSA). Injection volumes varied, depending on the protein amounts, but were typically 4.5μl. Samples were analysed by a nanoaquity LC system (Waters), interfaced to a Synapt G2S; operated in HDMSE (ion mobility) mode. Each sample was loaded on the Synapt G2S; and analysed concurrently. The HDMSE data was processed and searched using the Protein Lynx Global Server (PLGS), version 3, and protein identifications made using the Uniprot database (human reviewed). Scaffold (Proteome Software, Portland, Oregon, USA) was used to validate protein identifications derived from MS/MS sequencing results, visualised using the ProteinProphet computer algorithms (Searle, 2010). In Scaffold, the results from each individual paired gel slice sample were collated to give total protein identifications from each gel lane. Label-free quantitation was carried out using the “Top 3” method as described (Silva et al., 2006). In brief, the abundances of the three most abundant peptides for each 73 protein were compared to the abundances of a known amount of ADH (80 fmol) included in each sample analysed. Quantified amounts (fmol) for each identified protein, in each sample (each paired slice for shotgun method) were generated by PLGS and exported as csv files. Using a programme written in-house by Dr Ian Powley, the total amounts of each protein of interest for the whole sample were extracted.

### TRAIL/CD95 DISC Isolation

Isolation of the TRAIL/CD95 DISC was performed using a modified version of previously described method (Hughes et al., 2015). Briefly, 5 x 10^6^ cells were treated with 500 ng/ml of soluble biotinylated TRAIL (bTRAIL) or biotin labelled mAb to APO-1/Fas/CD95 (clone APO-1-3; Bender Medsystems; 400 ng/ml) and Protein A (50 ng/ml) at 4°C followed by 30 min at 37°C. Cells were incubated on ice for 1 h followed by either 20 (BJAB) or 25 (Jurkat) min at 37 °C. Cell pellets were washed and lysed on ice for 30 min. Lysates were cleared by centrifugation at 15,000 g for 30 min at 4 °C and the resulting supernatant incubated for 17 h at 4 °C on an end-to end rotator with 50 μl magnetic M-280 streptavidin Dynabeads (Dynal, Invitrogen). Beads were isolated from the supernatant and a sample of the supernatant was mixed with 10x sample buffer (0.5 M Tris-HCl pH 6.8, 0.4 % w/v bromophenol blue, 15 % v/v glycerol, 16 % w/v SDS, 5 % v/v β-mercaptoethanol). The beads were washed prior to the elution of proteins, which was achieved by boiling in 1x sample buffer. Proteins in the supernatants and bead eluates were separated by SDS-PAGE for analysis by Western blot.

### Separation of TRAIL DISC by sucrose density gradient centrifugation

Cells (100 x 10^6^) were treated as described above to induce DISC formation. The washed cell pellet was lysed in DISC lysis buffer and the cleared lysate loaded onto a continuous 10 - 45 % sucrose gradient (10 - 45 % w/v sucrose, 50 mM Tris-HCl pH 7.4, 150 mM NaCl, 0.1 % v/v Triton X-100) and centrifuged at 180,000 g for 17 h at 4 °C. Following centrifugation the gradients were fractionated into 0.5 ml samples. To each of the fractions 16 μl of magnetic M-280 streptavidin Dynabeads were added before overnight incubation at 4 °C. The beads were washed and the proteins eluted by boiling in 1x sample buffer for SDS-PAGE separation. Protein standards from high and low molecular weight kits were used as molecular weight markers for the gradient fractionation.

### Size-exclusion chromatography

Gel filtration column, Superdex 200 PC 3.2/30 (GE Healthcare), was used in conjunction with the SMART system (Pharmacia Biotech, now GE Healthcare). The running buffer was composed of 20 mM HEPES-KOH pH 7.0, 5 % w/v sucrose, 0.1 % w/v CHAPS and 5 mM dithiothretol (DTT). To this, 150 mM NaCl was added when using the Superdex 200 column. Samples were applied to the SMART system and eluted in 50 μl fractions at a rate of 40μl/min. The elution fractions were analysed though SDS-PAGE separation and Western blot. All columns were calibrated using protein standards from both high and low molecular weight kits (GE Healthcare).

### Flow cytometry analysis

Detection of Annexin V/Draq7 was used as a marker for apoptotic cells. 10,000 cells were analysed per sample, three biological repeats were performed per experiment and statistical differences determined by t-test.

Detection of cell surface expression of CD95: BJAB cells (1×10^6^) were re-suspended in 10 % v/v goat serum in PBS and incubated with 5 ng/μl of mouse monocolonal anti-Fas antibody (Clone CH-11) for 1 h on ice. Cells were washed and re-suspended in fresh 10 % goat serum. Fluorescein isothiocyanate (FITC)-conjugated goat anti-mouse immunoglobulin (F(ab’)_2_ fraction, Dako) was added to a final concentration of 50 ng/ml. Following incubation on ice for 1h, cells were washed and the level of CD95 receptor expression analysed using the BD FACS-Canto with excitation/emission wavelengths of 488/525 nm.

### Immunoprecipitation of FADD

Cell lysates prepared in lysis buffer were incubated at 4°C overnight with protein A/G beads and 1μg FADD antibody. Beads were pelleted and washed 3 times in lysis buffer. Precipitated proteins were eluted from the beads in 2x sample buffer. Resultant samples were analysed by SDS-PAGE/western blotting.

### RNA Interference

siRNA SMARTpools from Dharmacon were transiently transfected into BJAB cells by nucleofection, using the Amaxa Nucleofector II (Solution V, program C-009) according to the manufacturer’s instructions. The extent of protein knockdown was monitored by western blot analysis of cells 24, 48 and 72 h post-transfection.

**Table 1:** Antibodies used for western blot analysis.

| **Protein** | **Size**<br>**(approx, kDa)** | **Type** | **Company** | **Dilution** |
| --- | --- | --- | --- | --- |
| α-tubulin | 60 | Mouse monoclonal (DM1A) | Oncogene Research Products, Beeston, UK | 1:1000 |
| Bak | 30 | Rabbit polyclonal (NT) | Upstate, Millipore, Watford, UK | 1:1000 |
| Bcl-2 | 26 | Mouse monoclonal (124) | Dako, Ely, UK | 1:2000 |
| Bid | 22, 15, 13 | Rabbit polyclonal | Kind gift from Dr X Wang, University of Texas, USA | 1:2000 |
| Caspase-9 | 45, 35/37 | Mouse monoclonal (5B4) | Medical & Biological Laboratories, Woburn, USA | 1:1000 |
| c-FLIP | 55, 43, 26 | Rabbit polyclonal | Merck Frost Centre (Rasper et al, 1998) | 1:3000 |
| FADD | 24 | Mouse monoclonal (1) | BD Transduction Laboratories, Oxford, UK | 1:250 |
| CD95 | 48 | Rabbit polyclonal (C20) | Santa Cruz Biotechnology | 1:200 |
| MYLK2 | 65 | Rabbit polyclonal | Abcam | 1:1000 |
| Transferrin receptor | 85 | Mouse monoclonal (2) | BD Transduction Laboratories | 1:1000 |
| TRAIL-R1 | 57 | Rabbit polyclonal (CT) | Prosci Incorporated, Poway, USA | 1:1000 |
| TRAIL-R2 | 48, 40 | Rabbit polyclonal | Cell Signaling Technology | 1:1000 |
| XIAP | 57 | Mouse monoclonal (28) | BD Transduction Laboratories | 1:2000 |
| Caspase-8 | 55/53, 43,41 | Mouse monoclonal (C15) | Adipogen Life Sciences | 1:500 |
| Caspase-3 | 32, 19/17 | Rabbit polyclonal | Kind gift from Dr X-M Sun, MRC Toxicology Unit | 1:2000 |
| Caspase-7 | 34, 19 | Rabbit polyclonal | Kind gift from Dr X-M Sun, MRC Toxicology Unit | 1:2000 |
| Actin | 42 | Mouse monoclonal (AC-15) | Sigma | 1:4000 |

## References

Alappat, E.C., Feig, C., Boyerinas, B., Volkland, J., Samuels, M., Murmann, A.E., Thorburn, A., Kidd, V.J., Slaughter, C.A., Osborn, S.L., et al. (2005). Phosphorylation of FADD at serine 194 by CKIalpha regulates its nonapoptotic activities. Mol Cell 19, 321–332.

Algeciras-Schimnich, A., Shen, L., Barnhart, B.C., Murmann, A.E., Burkhardt, J.K., and Peter, M.E. (2002). Molecular ordering of the initial signaling events of CD95. Mol Cell Biol 22, 207–220.

Blanchard, H., Kodandapani, L., Mittl, P.R., Marco, S.D., Krebs, J.F., Wu, J.C., Tomaselli, K.J., and Grutter, M.G. (1999). The three-dimensional structure of caspase-8: an initiator enzyme in apoptosis. Structure 7, 1125–1133.

Boldin, M.P., Goncharov, T.M., Goltsev, Y.V., and Wallach, D. (1996). Involvement of MACH, a novel MORT1/FADD-interacting protease, in Fas/APO-1- and TNF receptor-induced cell death. Cell 85, 803–815.

Chinnaiyan, A.M., O’Rourke, K., Tewari, M., and Dixit, V.M. (1995). FADD, a novel death domain-containing protein, interacts with the death domain of Fas and initiates apoptosis. Cell 81, 505–512.

Clancy, L., Mruk, K., Archer, K., Woelfel, M., Mongkolsapaya, J., Screaton, G., Lenardo, M.J., and Chan, F.K. (2005). Preligand assembly domain-mediated ligand-independent association between TRAIL receptor 4 (TR4) and TR2 regulates TRAIL-induced apoptosis. Proc Natl Acad Sci U S A 102, 18099–18104.

Dickens, L.S., Boyd, R.S., Jukes-Jones, R., Hughes, M.A., Robinson, G.L., Fairall, L., Schwabe, J.W., Cain, K., and Macfarlane, M. (2012). A death effector domain chain DISC model reveals a crucial role for caspase-8 chain assembly in mediating apoptotic cell death. Mol Cell 47, 291–305.

Dudek, S.M., Chiang, E.T., Camp, S.M., Guo, Y., Zhao, J., Brown, M.E., Singleton, P.A., Wang, L., Desai, A., Arce, F.T., et al. (2010). Abl tyrosine kinase phosphorylates nonmuscle Myosin light chain kinase to regulate endothelial barrier function. Mol Biol Cell 21, 4042–4056.

Esposito, D., Sankar, A., Morgner, N., Robinson, C.V., Rittinger, K., and Driscoll, P.C. (2010). Solution NMR investigation of the CD95/FADD homotypic death domain complex suggests lack of engagement of the CD95 C terminus. Structure 18, 1378–1390.

Fassl, S., Leisser, C., Huettenbrenner, S., Maier, S., Rosenberger, G., Strasser, S., Grusch, M., Fuhrmann, G., Leuhuber, K., Polgar, D., et al. (2003). Transferrin ensures survival of ovarian carcinoma cells when apoptosis is induced by TNFalpha, FasL, TRAIL, or Myc. Oncogene 22, 8343–8355.

Fox, J.L., Hughes, M.A., Meng, X., Sarnowska, N.A., Powley, I.R., Jukes-Jones, R., Dinsdale, D., Ragan, T.J., Fairall, L., Schwabe, J.W.R., et al. (2021). Cryo-EM structural analysis of FADD:Caspase-8 complexes defines the catalytic dimer architecture for co-ordinated control of cell fate. Nat Commun 12, 819.

Fu, T.M., Li, Y., Lu, A., Li, Z., Vajjhala, P.R., Cruz, A.C., Srivastava, D.B., DiMaio, F., Penczek, P.A., Siegel, R.M., et al. (2016). Cryo-EM Structure of Caspase-8 Tandem DED Filament Reveals Assembly and Regulation Mechanisms of the Death-Inducing Signaling Complex. Mol Cell 64, 236–250.

Gomez-Angelats, M., and Cidlowski, J.A. (2003). Molecular evidence for the nuclear localization of FADD. Cell Death Differ 10, 791–797.

Habashy, H.O., Powe, D.G., Staka, C.M., Rakha, E.A., Ball, G., Green, A.R., Aleskandarany, M., Paish, E.C., Douglas Macmillan, R., Nicholson, R.I., et al. (2010). Transferrin receptor (CD71) is a marker of poor prognosis in breast cancer and can predict response to tamoxifen. Breast Cancer Res Treat 119, 283–293.

Huang, B., Eberstadt, M., Olejniczak, E.T., Meadows, R.P., and Fesik, S.W. (1996). NMR structure and mutagenesis of the Fas (APO-1/CD95) death domain. Nature 384, 638–641.

Hughes, M.A., Langlais, C., Cain, K., and MacFarlane, M. (2015). Activation, Isolation, and Analysis of the Death-Inducing Signaling Complex. Cold Spring Harb Protoc 2015, pdb prot087098.

Hughes, M.A., Powley, I.R., Jukes-Jones, R., Horn, S., Feoktistova, M., Fairall, L., Schwabe, J.W., Leverkus, M., Cain, K., and MacFarlane, M. (2016). Co-operative and Hierarchical Binding of c-FLIP and Caspase-8: A Unified Model Defines How c-FLIP Isoforms Differentially Control Cell Fate. Mol Cell 61, 834–849.

Jin, Y., Atkinson, S.J., Marrs, J.A., and Gallagher, P.J. (2001). Myosin ii light chain phosphorylation regulates membrane localization and apoptotic signaling of tumor necrosis factor receptor-1. J Biol Chem 276, 30342–30349.

Juo, P., Woo, M.S., Kuo, C.J., Signorelli, P., Biemann, H.P., Hannun, Y.A., and Blenis, J. (1999). FADD is required for multiple signaling events downstream of the receptor Fas. Cell Growth Differ 10, 797–804.

Kasibhatla, S., Jessen, K.A., Maliartchouk, S., Wang, J.Y., English, N.M., Drewe, J., Qiu, L., Archer, S.P., Ponce, A.E., Sirisoma, N., et al. (2005). A role for transferrin receptor in triggering apoptosis when targeted with gambogic acid. Proc Natl Acad Sci U S A 102, 12095–12100.

Keller, N., Mares, J., Zerbe, O., and Grutter, M.G. (2009). Structural and biochemical studies on procaspase-8: new insights on initiator caspase activation. Structure 17, 438–448.

Khadra, N., Bresson-Bepoldin, L., Penna, A., Chaigne-Delalande, B., Segui, B., Levade, T., Vacher, A.M., Reiffers, J., Ducret, T., Moreau, J.F., et al. (2011). CD95 triggers Orai1-mediated localized Ca2+ entry, regulates recruitment of protein kinase C (PKC) beta2, and prevents death-inducing signaling complex formation. Proc Natl Acad Sci U S A 108, 19072–19077.

Kim, D.Y., and Helfman, D.M. (2016). Loss of MLCK leads to disruption of cell-cell adhesion and invasive behavior of breast epithelial cells via increased expression of EGFR and ERK/JNK signaling. Oncogene 35, 4495–4508.

Kischkel, F.C., Hellbardt, S., Behrmann, I., Germer, M., Pawlita, M., Krammer, P.H., and Peter, M.E. (1995). Cytotoxicity-dependent APO-1 (Fas/CD95)-associated proteins form a death-inducing signaling complex (DISC) with the receptor. Embo J 14, 5579–5588.

Lee, E.W., Kim, J.H., Ahn, Y.H., Seo, J., Ko, A., Jeong, M., Kim, S.J., Ro, J.Y., Park, K.M., Lee, H.W., et al. (2012). Ubiquitination and degradation of the FADD adaptor protein regulate death receptor-mediated apoptosis and necroptosis. Nat Commun 3, 978.

Martin, D.A., Zheng, L., Siegel, R.M., Huang, B., Fisher, G.H., Wang, J., Jackson, C.E., Puck, J.M., Dale, J., Straus, S.E., et al. (1999). Defective CD95/APO-1/Fas signal complex formation in the human autoimmune lymphoproliferative syndrome, type Ia. Proc Natl Acad Sci U S A 96, 4552–4557.

Naismith, J.H., and Sprang, S.R. (1998). Modularity in the TNF-receptor family. Trends Biochem Sci 23, 74–79.

Newton, K., Harris, A.W., Bath, M.L., Smith, K.G., and Strasser, A. (1998). A dominant interfering mutant of FADD/MORT1 enhances deletion of autoreactive thymocytes and inhibits proliferation of mature T lymphocytes. Embo J 17, 706–718.

Newton, K., Kurts, C., Harris, A.W., and Strasser, A. (2001). Effects of a dominant interfering mutant of FADD on signal transduction in activated T cells. Curr Biol 11, 273–276.

Parsons, D.W., Wang, T.L., Samuels, Y., Bardelli, A., Cummins, J.M., DeLong, L., Silliman, N., Ptak, J., Szabo, S., Willson, J.K., et al. (2005). Colorectal cancer: mutations in a signalling pathway. Nature 436, 792.

Pop, C., Fitzgerald, P., Green, D.R., and Salvesen, G.S. (2007). Role of proteolysis in caspase-8 activation and stabilization. Biochemistry 46, 4398–4407.

Rosager, A.M., Sorensen, M.D., Dahlrot, R.H., Hansen, S., Schonberg, D.L., Rich, J.N., Lathia, J.D., and Kristensen, B.W. (2017). Transferrin receptor-1 and ferritin heavy and light chains in astrocytic brain tumors: Expression and prognostic value. PLoS One 12, e0182954.

Schleich, K., Warnken, U., Fricker, N., Ozturk, S., Richter, P., Kammerer, K., Schnolzer, M., Krammer, P.H., and Lavrik, I.N. (2012). Stoichiometry of the CD95 death-inducing signaling complex: experimental and modeling evidence for a death effector domain chain model. Mol Cell 47, 306–319.

Scott, F.L., Stec, B., Pop, C., Dobaczewska, M.K., Lee, J.J., Monosov, E., Robinson, H., Salvesen, G.S., Schwarzenbacher, R., and Riedl, S.J. (2009). The Fas-FADD death domain complex structure unravels signalling by receptor clustering. Nature 457, 1019–1022.

Searle, B.C. (2010). Scaffold: a bioinformatic tool for validating MS/MS-based proteomic studies. Proteomics 10, 1265–1269.

Siegel, R.M., Martin, D.A., Zheng, L., Ng, S.Y., Bertin, J., Cohen, J., and Lenardo, M.J. (1998). Death-effector filaments: novel cytoplasmic structures that recruit caspases and trigger apoptosis. J Cell Biol 141, 1243–1253.

Siegel, R.M., Muppidi, J.R., Sarker, M., Lobito, A., Jen, M., Martin, D., Straus, S.E., and Lenardo, M.J. (2004). SPOTS: signaling protein oligomeric transduction structures are early mediators of death receptor-induced apoptosis at the plasma membrane. J Cell Biol 167, 735–744.

Silva, J.C., Gorenstein, M.V., Li, G.Z., Vissers, J.P., and Geromanos, S.J. (2006). Absolute quantification of proteins by LCMSE: a virtue of parallel MS acquisition. Mol Cell Proteomics 5, 144–156.

Stull, J.T., Kamm, K.E., and Vandenboom, R. (2011). Myosin light chain kinase and the role of myosin light chain phosphorylation in skeletal muscle. Arch Biochem Biophys 510, 120–128.

Tourneur, L., and Chiocchia, G. (2010). FADD: a regulator of life and death. Trends Immunol 31, 260–269.

Usatyuk, P.V., Singleton, P.A., Pendyala, S., Kalari, S.K., He, D., Gorshkova, I.A., Camp, S.M., Moitra, J., Dudek, S.M., Garcia, J.G., et al. (2012). Novel role for non-muscle myosin light chain kinase (MLCK) in hyperoxia-induced recruitment of cytoskeletal proteins, NADPH oxidase activation, and reactive oxygen species generation in lung endothelium. J Biol Chem 287, 9360–9375.

Vilmont, V., Filhol, O., Hesse, A.M., Coute, Y., Hue, C., Remy-Tourneur, L., Mistou, S., Cochet, C., and Chiocchia, G. (2015). Modulatory role of the anti-apoptotic protein kinase CK2 in the sub-cellular localization of Fas associated death domain protein (FADD). Biochim Biophys Acta 1853, 2885–2896.

Wang, L., Yang, J.K., Kabaleeswaran, V., Rice, A.J., Cruz, A.C., Park, A.Y., Yin, Q., Damko, E., Jang, S.B., Raunser, S., et al. (2010a). The Fas-FADD death domain complex structure reveals the basis of DISC assembly and disease mutations. Nat Struct Mol Biol 17, 1324–1329.

Wang, Z., Watt, W., Brooks, N.A., Harris, M.S., Urban, J., Boatman, D., McMillan, M., Kahn, M., Heinrikson, R.L., Finzel, B.C., et al. (2010b). Kinetic and structural characterization of caspase-3 and caspase-8 inhibition by a novel class of irreversible inhibitors. Biochim Biophys Acta 1804, 1817–1831.

Watt, W., Koeplinger, K.A., Mildner, A.M., Heinrikson, R.L., Tomasselli, A.G., and Watenpaugh, K.D. (1999). The atomic-resolution structure of human caspase-8, a key activator of apoptosis. Structure 7, 1135–1143.

Werner, M.H., Wu, C., and Walsh, C.M. (2006). Emerging roles for the death adaptor FADD in death receptor avidity and cell cycle regulation. Cell Cycle 5, 2332–2338.

Wozniak, A.L., Wang, X., Stieren, E.S., Scarbrough, S.G., Elferink, C.J., and Boehning, D. (2006). Requirement of biphasic calcium release from the endoplasmic reticulum for Fas-mediated apoptosis. J Cell Biol 175, 709–714.

