## Supplementary Data File for "Characterisation of FADD interactome reveals novel insights into FADD recruitment and signalling at the Death Inducing Signalling Complex (DISC)"

| Protein ID | Fold Change |
|---|---|
| LAC2_HUMAN | 1 |
| KV302_HUMAN | 411.69 |
| LAC7_HUMAN | 1 |
| LAC6_HUMAN | 0.05 |
| KV305_HUMAN | 1 |
| KV304_HUMAN | 1.58 |
| KV307_HUMAN | 1.31 |
| IGKC_HUMAN | 0.9 |
| KV303_HUMAN | 141.01 |
| IGLL5_HUMAN | 0.79 |
| ADH1_YEAST | 1 |
| KV309_HUMAN | 1 |
| HV303_HUMAN | 1.96 |
| LV302_HUMAN | 1.32 |
| HV316_HUMAN | 0.04 |
| HV305_HUMAN | 1 |
| TBB3_HUMAN | 0.08 |
| KV206_HUMAN | 1.33 |
| HS90B_HUMAN | 0.87 |
| H4_HUMAN | 1.37 |
| PROF1_HUMAN | 0.01 |
| ALBU_BOVIN | 1120.89 |
| HS90A_HUMAN | 0.99 |
| TBA1B_HUMAN | 1 |
| TBA1C_HUMAN | 1.47 |
| ACTB_HUMAN | 0.01 |
| TBA1A_HUMAN | 1 |
| LV106_HUMAN | 0.01 |
| KV402_HUMAN | 1.46 |
| LV102_HUMAN | 0.9 |
| KV204_HUMAN | 1 |
| G3P_HUMAN | 0.01 |
| HV304_HUMAN | 1 |
| TBA3C_HUMAN | 1 |
| ACTA_HUMAN | 1 |
| IGHG1_HUMAN | 1.36 |
| HSP7C_HUMAN | 0.95 |
| HS902_HUMAN | 1 |
| KV118_HUMAN | 1.62 |
| KV311_HUMAN | 0.02 |
| TCPQ_HUMAN | 0.01 |
| TBB5_HUMAN | 0.91 |
| H2B1N_HUMAN | 1 |
| IGHG2_HUMAN | 1.26 |
| H90B3_HUMAN | 1 |
| POTEJ_HUMAN | 1 |
| RS20_HUMAN | 1.53 |
| SUMO2_HUMAN | 1 |
| RL40_HUMAN | 0.88 |
| RS16_HUMAN | 1.19 |
| PEBB_HUMAN | 1 |
| RO52_HUMAN | 0.01 |
| H2B1C_HUMAN | 0.02 |
| LDHA_HUMAN | 0.82 |
| K2C1_HUMAN | 0.71 |
| H2B1B_HUMAN | 0.03 |
| LDHB_HUMAN | 0.88 |

| | |
|---|---|
| TBB2A_HUMAN | 1.38 |
| EF1A1_HUMAN | 0.01 |
| IGHG4_HUMAN | 0.95 |
| RSSA_HUMAN | 1.07 |
| KV107_HUMAN | 1 |
| KV405_HUMAN | 0.03 |
| LV301_HUMAN | 1.31 |
| KPYM_HUMAN | 0.82 |
| H31T_HUMAN | 1.09 |
| FADD_HUMAN | 0.46 |
| ACTC_HUMAN | 1 |
| TBA4A_HUMAN | 0.03 |
| POTEE_HUMAN | 0.03 |
| KV123_HUMAN | 1 |
| ACTH_HUMAN | 31.8 |
| ATPA_HUMAN | 0.96 |
| KV310_HUMAN | 1 |
| KV110_HUMAN | 1.29 |
| NUCL_HUMAN | 0.01 |
| TCPZ_HUMAN | 0.01 |
| K1C10_HUMAN | 0.64 |
| FKB1A_HUMAN | 1 |
| PHB2_HUMAN | 0.03 |
| CH60_HUMAN | 0.6 |
| EF1D_HUMAN | 0.87 |
| KV116_HUMAN | 1.6 |
| KV401_HUMAN | 1 |
| KV122_HUMAN | 1 |
| RS3_HUMAN | 1.23 |
| RLA0_HUMAN | 0.03 |
| TCPD_HUMAN | 0.01 |
| HS71A_HUMAN | 0.01 |
| RS18_HUMAN | 1.15 |
| RL22_HUMAN | 0.82 |
| ACTBM_HUMAN | 24.02 |
| IF4A2_HUMAN | 1 |
| SRP14_HUMAN | 0.92 |
| CH10_HUMAN | 31.17 |
| MYL6_HUMAN | 0.91 |
| KV117_HUMAN | 0.21 |
| KV115_HUMAN | 0.16 |
| KV102_HUMAN | 1 |
| KV101_HUMAN | 1 |
| IGHG3_HUMAN | 1.37 |
| HSP72_HUMAN | 1 |
| TCPB_HUMAN | 0.01 |
| ENPL_HUMAN | 0.8 |
| TERA_HUMAN | 0.95 |
| TCPA_HUMAN | 0.66 |
| ENOA_HUMAN | 1.01 |
| KV306_HUMAN | 1 |
| RS19_HUMAN | 1.12 |
| RS7_HUMAN | 1.06 |
| GNAS2_HUMAN | 1 |
| HV320_HUMAN | 1.65 |
| TCPG_HUMAN | 0.86 |
| F10A1_HUMAN | 1 |
| LV403_HUMAN | 1 |
| RS14_HUMAN | 0.84 |
| ACTBL_HUMAN | 0.83 |

| | |
|---|---|
| TBB6_HUMAN | 0.45 |
| H2A1B_HUMAN | 1 |
| COR1A_HUMAN | 1.09 |
| SERA_HUMAN | 0.03 |
| K22E_HUMAN | 2.97 |
| GRP78_HUMAN | 0.97 |
| LV104_HUMAN | 1 |
| POTEF_HUMAN | 1 |
| PRDX1_HUMAN | 0.66 |
| ST134_HUMAN | 0.06 |
| RLA2_HUMAN | 0.8 |
| H2AV_HUMAN | 0.08 |
| RL27_HUMAN | 0.7 |
| SFPQ_HUMAN | 1.08 |
| TBA4B_HUMAN | 1 |
| PRDX2_HUMAN | 0.05 |
| EF1B_HUMAN | 0.52 |
| H2A1H_HUMAN | 1.24 |
| ENOB_HUMAN | 1 |
| EF1G_HUMAN | 0.02 |
| TFR1_HUMAN | 1.2 |
| TCPH_HUMAN | 0.91 |
| TBB4A_HUMAN | 0.05 |
| ML12A_HUMAN | 0.76 |
| ATPB_HUMAN | 0.89 |
| YBOX1_HUMAN | 0.81 |
| KV113_HUMAN | 3.85 |
| PHB_HUMAN | 0.91 |
| EF1A2_HUMAN | 14.47 |
| MOES_HUMAN | 0.02 |
| RS25_HUMAN | 1 |
| PGAM4_HUMAN | 1 |
| ATD3A_HUMAN | 0.84 |
| XRCC5_HUMAN | 0.04 |
| TBB4B_HUMAN | 1 |
| H90B2_HUMAN | 1.01 |
| STMN1_HUMAN | 0.82 |
| HNRH1_HUMAN | 0.94 |
| XRCC6_HUMAN | 0.03 |
| RS13_HUMAN | 1.18 |
| LV001_HUMAN | 1.25 |
| HNRPK_HUMAN | 0.75 |
| RS23_HUMAN | 0.07 |
| ARPC4_HUMAN | 1.05 |
| ADT2_HUMAN | 1 |
| PLSL_HUMAN | 0.82 |
| TCPE_HUMAN | 0.83 |
| PIMT_HUMAN | 0.2 |
| 1433Z_HUMAN | 1.18 |
| CUX1_HUMAN | 1 |
| TRYP_PIG | 598.75 |
| RBBP4_HUMAN | 1.03 |
| IF4H_HUMAN | 1 |
| K1C9_HUMAN | 0.7 |
| COF1_HUMAN | 0.06 |
| RL35_HUMAN | 0.09 |
| ANXA1_HUMAN | 1.02 |
| EZRI_HUMAN | 10.81 |
| GRP75_HUMAN | 0.02 |
| RO60_HUMAN | 0.02 |

| | |
|---|---|
| UBC9_HUMAN | 1 |
| AIMP1_HUMAN | 0.05 |
| ATD3B_HUMAN | 1.46 |
| RBBP7_HUMAN | 1 |
| RL31_HUMAN | 0.95 |
| PSMD3_HUMAN | 0.85 |
| 1433E_HUMAN | 1.02 |
| PGK1_HUMAN | 0.02 |
| TRAP1_HUMAN | 0.26 |
| RCN2_HUMAN | 0.03 |
| RL7A_HUMAN | 0.06 |
| MCM7_HUMAN | 1.49 |
| YBOX3_HUMAN | 5.55 |
| ADT3_HUMAN | 0.93 |
| K2C1B_HUMAN | 0.95 |
| IMDH2_HUMAN | 0.84 |
| NUDC_HUMAN | 0.04 |
| THIO_HUMAN | 33.56 |
| PPAC_HUMAN | 0.82 |
| NONO_HUMAN | 0.84 |
| CSK21_HUMAN | 0.04 |
| CN166_HUMAN | 0.99 |
| C1QBP_HUMAN | 1.37 |
| K2C6B_HUMAN | 0.56 |
| ROA2_HUMAN | 0.84 |
| RS2_HUMAN | 0.99 |
| PRS8_HUMAN | 1.01 |
| RL7_HUMAN | 0.72 |
| KV308_HUMAN | 1 |
| RL12_HUMAN | 0.21 |
| RLA0L_HUMAN | 1 |
| K2C7_HUMAN | 1 |
| SET_HUMAN | 0.05 |
| DNJA1_HUMAN | 0.83 |
| RS15A_HUMAN | 1.7 |
| F10A5_HUMAN | 1 |
| FUBP2_HUMAN | 0.84 |
| ANM1_HUMAN | 0.06 |
| HNRPF_HUMAN | 0.03 |
| SYDC_HUMAN | 0.04 |
| MCM6_HUMAN | 1.23 |
| RL11_HUMAN | 0.76 |
| CAZA1_HUMAN | 0.81 |
| PPIA_HUMAN | 0.86 |
| CALM_HUMAN | 0.86 |
| IF4A1_HUMAN | 0.04 |
| PRDX3_HUMAN | 0.87 |
| PAXX_HUMAN | 1.01 |
| BC11B_HUMAN | 1 |
| MDHC_HUMAN | 0.69 |
| EF2_HUMAN | 0.76 |
| H2A2B_HUMAN | 1 |
| RS3A_HUMAN | 1.09 |
| RAB1C_HUMAN | 1 |
| PSB2_HUMAN | 0.03 |
| PSA2_HUMAN | 0.06 |
| RL36L_HUMAN | 1 |
| IGLL1_HUMAN | 45 |
| MTPN_HUMAN | 1 |
| STIP1_HUMAN | 0.04 |

| | |
|---|---|
| PSMD2_HUMAN | 0.84 |
| ATPD_HUMAN | 0.06 |
| DDX1_HUMAN | 0.84 |
| KV104_HUMAN | 0.69 |
| RS10_HUMAN | 1.27 |
| RL17_HUMAN | 0.94 |
| PP2AA_HUMAN | 1.19 |
| MGN_HUMAN | 1 |
| PRDX4_HUMAN | 2.35 |
| SSBP_HUMAN | 1.16 |
| PDIA6_HUMAN | 0.05 |
| RS4X_HUMAN | 1.2 |
| RS28_HUMAN | 1 |
| LV103_HUMAN | 2.23 |
| CSK23_HUMAN | 1 |
| RS9_HUMAN | 0.75 |
| CDK1_HUMAN | 0.56 |
| HV103_HUMAN | 0.04 |
| MCM3_HUMAN | 1.01 |
| DDX5_HUMAN | 0.61 |
| RL27A_HUMAN | 0.7 |
| NACA_HUMAN | 1 |
| ALDOA_HUMAN | 0.91 |
| PDCD5_HUMAN | 0.1 |
| RPN1_HUMAN | 0.05 |
| RS26_HUMAN | 1.1 |
| MCM5_HUMAN | 0.77 |
| SYQ_HUMAN | 0.09 |
| DJB11_HUMAN | 0.89 |
| LAP2B_HUMAN | 10.59 |
| PSB3_HUMAN | 0.92 |
| NPM_HUMAN | 40.57 |
| PUR6_HUMAN | 1.38 |
| KV301_HUMAN | 130.68 |
| HSP74_HUMAN | 0.44 |
| ACTN1_HUMAN | 0.06 |
| PSD11_HUMAN | 0.04 |
| ACTN4_HUMAN | 12.31 |
| PSA4_HUMAN | 0.05 |
| PGK2_HUMAN | 1 |
| PDIA3_HUMAN | 0.08 |
| IF5A2_HUMAN | 1 |
| ODPB_HUMAN | 0.06 |
| NDKB_HUMAN | 1 |
| TALDO_HUMAN | 1 |
| LV206_HUMAN | 1 |
| HNRPM_HUMAN | 0.71 |
| LANC1_HUMAN | 0.08 |
| COF2_HUMAN | 14.95 |
| PSA5_HUMAN | 1.45 |
| MYH9_HUMAN | 49.86 |
| 6PGL_HUMAN | 1 |
| PRS7_HUMAN | 0.95 |
| TF65_HUMAN | 1 |
| DHX36_HUMAN | 0.61 |
| TPM3_HUMAN | 1.05 |
| ERH_HUMAN | 1 |
| 6PGD_HUMAN | 0.75 |
| ALBU_HUMAN | 42.57 |
| PSA7_HUMAN | 1.29 |

| | |
|---|---|
| PRP19_HUMAN | 0.83 |
| RAB8A_HUMAN | 1 |
| HNRPC_HUMAN | 0.06 |
| RS6_HUMAN | 0.97 |
| VIME_HUMAN | 0.06 |
| VRK3_HUMAN | 1 |
| SMD3_HUMAN | 1 |
| LA_HUMAN | 0.97 |
| RTCB_HUMAN | 1.33 |
| HV306_HUMAN | 1 |
| CYTB_HUMAN | 1 |
| NDKA_HUMAN | 0.77 |
| MAT2B_HUMAN | 0.09 |
| CAP1_HUMAN | 106.49 |
| CNPY2_HUMAN | 1 |
| HS105_HUMAN | 0.06 |
| K1C17_HUMAN | 0.18 |
| ANXA6_HUMAN | 0.07 |
| RS12_HUMAN | 16.91 |
| RL8_HUMAN | 0.8 |
| FUMH_HUMAN | 0.8 |
| IMB1_HUMAN | 0.85 |
| PSDE_HUMAN | 0.89 |
| LAP2A_HUMAN | 12.43 |
| 1B14_HUMAN | 1 |
| IF5A1_HUMAN | 1 |
| RL10A_HUMAN | 0.96 |
| KPYR_HUMAN | 1 |
| PNPH_HUMAN | 0.93 |
| HMBX1_HUMAN | 1 |
| ACLY_HUMAN | 0.02 |
| SRSF2_HUMAN | 0.31 |
| ARF1_HUMAN | 0.34 |
| ATPG_HUMAN | 0.79 |
| RL24_HUMAN | 1.18 |
| SSRP1_HUMAN | 0.88 |
| RL18_HUMAN | 8.55 |
| RL4_HUMAN | 0.82 |
| 4F2_HUMAN | 0.09 |
| 1433B_HUMAN | 1.18 |
| ROA1_HUMAN | 0.88 |
| MCM2_HUMAN | 0.06 |
| KPRB_HUMAN | 0.83 |
| H12_HUMAN | 0.99 |
| CAZA2_HUMAN | 0.13 |
| K2C4_HUMAN | 1 |
| SUH_HUMAN | 0.06 |
| MCM4_HUMAN | 0.05 |
| HSP77_HUMAN | 1 |
| K22O_HUMAN | 0.02 |
| FLNA_HUMAN | 1.37 |
| NACAM_HUMAN | 0.12 |
| GLYM_HUMAN | 0.79 |
| PSB5_HUMAN | 0.43 |
| RUVB2_HUMAN | 1.42 |
| PRS6B_HUMAN | 1.01 |
| HINT1_HUMAN | 1 |
| LV209_HUMAN | 1 |
| HNRPU_HUMAN | 1 |
| LV501_HUMAN | 29.98 |

| | |
|---|---|
| CYC_HUMAN | 1 |
| PSPC1_HUMAN | 0.82 |
| MACF1_HUMAN | 1 |
| SYK_HUMAN | 0.04 |
| RS24_HUMAN | 5.98 |
| HCD2_HUMAN | 1 |
| PRS4_HUMAN | 0.05 |
| SEPT1_HUMAN | 0.07 |
| DPYL2_HUMAN | 0.98 |
| PLEC_HUMAN | 1 |
| DDX17_HUMAN | 0.05 |
| K1C27_HUMAN | 0.14 |
| PCNA_HUMAN | 0.69 |
| IF2B_HUMAN | 0.4 |
| RSU1_HUMAN | 0.71 |
| OST48_HUMAN | 1.54 |
| PSME3_HUMAN | 0.84 |
| METK2_HUMAN | 0.06 |
| SEPT7_HUMAN | 0.93 |
| PTPRC_HUMAN | 3.02 |
| GNAO_HUMAN | 1 |
| GDIB_HUMAN | 0.77 |
| ENOG_HUMAN | 0.08 |
| RL23_HUMAN | 3.72 |
| RAB35_HUMAN | 1 |
| SF3A3_HUMAN | 0.08 |
| GBLP_HUMAN | 0.88 |
| SORCN_HUMAN | 0.9 |
| PCBP2_HUMAN | 0.9 |
| PSD12_HUMAN | 0.86 |
| TIF1B_HUMAN | 0.79 |
| RL14_HUMAN | 0.06 |
| K1C24_HUMAN | 8.16 |
| RL28_HUMAN | 0.83 |
| PCBP1_HUMAN | 0.04 |
| ARP3_HUMAN | 0.14 |
| COPD_HUMAN | 0.77 |
| KV112_HUMAN | 1 |
| DIAP1_HUMAN | 0.02 |
| ATP5H_HUMAN | 0.38 |
| FA98B_HUMAN | 0.05 |
| SAHH_HUMAN | 0.91 |
| RL3_HUMAN | 0.84 |
| SYG_HUMAN | 0.82 |
| COPG1_HUMAN | 1.09 |
| K2C75_HUMAN | 3.22 |
| PSA1_HUMAN | 0.03 |
| 1433G_HUMAN | 1.62 |
| CDC42_HUMAN | 1 |
| SF3B6_HUMAN | 0.19 |
| K1C28_HUMAN | 1.63 |
| GBB1_HUMAN | 1 |
| CSN1_HUMAN | 0.75 |
| SYLC_HUMAN | 1.02 |
| NDK8_HUMAN | 0.66 |
| KAPCB_HUMAN | 1 |
| IF2G_HUMAN | 0.1 |
| IPYR_HUMAN | 1 |
| TBG1_HUMAN | 0.78 |
| SH3L1_HUMAN | 1 |

| | |
|---|---|
| DCAF7_HUMAN | 0.84 |
| HV206_HUMAN | 0.07 |
| H15_HUMAN | 0.79 |
| ARF3_HUMAN | 2.66 |
| SEPT6_HUMAN | 16.35 |
| RAC2_HUMAN | 0.57 |
| GAPR1_HUMAN | 1 |
| TFG_HUMAN | 0.85 |
| RL34_HUMAN | 0.83 |
| SEC13_HUMAN | 0.18 |
| ARPC5_HUMAN | 1 |
| CDK2_HUMAN | 0.62 |
| RUVB1_HUMAN | 0.06 |
| AT5F1_HUMAN | 1.51 |
| EFTU_HUMAN | 0.46 |
| 1433F_HUMAN | 1.29 |
| ODP2_HUMAN | 0.18 |
| 1433T_HUMAN | 1.64 |
| LV101_HUMAN | 0.01 |
| KINH_HUMAN | 0.13 |
| PTN6_HUMAN | 0.13 |
| PPR18_HUMAN | 1 |
| ELOB_HUMAN | 0.11 |
| RU2B_HUMAN | 0.15 |
| CLH1_HUMAN | 0.05 |
| RL15_HUMAN | 1.2 |
| TPIS_HUMAN | 0.08 |
| TOM34_HUMAN | 1 |
| THIC_HUMAN | 1 |
| SRSF1_HUMAN | 0.94 |
| G6PI_HUMAN | 0.03 |
| CISY_HUMAN | 1 |
| PRKDC_HUMAN | 1 |
| SF3B3_HUMAN | 0.74 |
| GANAB_HUMAN | 0.98 |
| EIF3F_HUMAN | 0.9 |
| LV212_HUMAN | 55.65 |
| G6PD_HUMAN | 0.08 |
| CPSF5_HUMAN | 1.3 |
| PARVG_HUMAN | 1.09 |
| 2AAA_HUMAN | 0.91 |
| DCD_HUMAN | 56.26 |
| DHX15_HUMAN | 0.42 |
| PFD5_HUMAN | 1 |
| CD2B2_HUMAN | 1 |
| COPB_HUMAN | 0.08 |
| SYIC_HUMAN | 0.07 |
| TPM4_HUMAN | 1.73 |
| SPTB2_HUMAN | 0.45 |
| LSM6_HUMAN | 1 |
| K2C78_HUMAN | 1 |
| RS8_HUMAN | 1.79 |
| COPA_HUMAN | 0.11 |
| RS30_HUMAN | 1 |
| NP1L1_HUMAN | 0.81 |
| PGAM5_HUMAN | 1.04 |
| RL23A_HUMAN | 0.98 |
| K2C5_HUMAN | 0.74 |
| C1TC_HUMAN | 0.84 |
| HS904_HUMAN | 1 |

| | |
|---|---|
| HDAC2_HUMAN | 0.22 |
| PPID_HUMAN | 1 |
| HV302_HUMAN | 0.05 |
| 1433S_HUMAN | 0.04 |
| RAB10_HUMAN | 1 |
| GRAP2_HUMAN | 0.99 |
| HDAC1_HUMAN | 0.63 |
| MRP_HUMAN | 0.38 |
| SEPT2_HUMAN | 0.13 |
| GDIR2_HUMAN | 0.9 |
| SEPT9_HUMAN | 0.04 |
| RHOG_HUMAN | 1 |
| RBM39_HUMAN | 0.29 |
| DAD1_HUMAN | 7.85 |
| KAP0_HUMAN | 0.06 |
| UBA1_HUMAN | 0.07 |
| ATP5I_HUMAN | 9.08 |
| KV120_HUMAN | 1.39 |
| YTHD3_HUMAN | 0.2 |
| RL35A_HUMAN | 0.19 |
| LYRIC_HUMAN | 0.47 |
| SF3A1_HUMAN | 0.84 |
| PHF5A_HUMAN | 6.42 |
| MPCP_HUMAN | 0.92 |
| RAB8B_HUMAN | 0 |
| AMPL_HUMAN | 0.12 |
| RT27_HUMAN | 0.77 |
| RM01_HUMAN | 0.78 |
| CHIP_HUMAN | 0.79 |
| RL18A_HUMAN | 0.27 |
| SRSF5_HUMAN | 0.1 |
| GFPT1_HUMAN | 0.12 |
| CDK7_HUMAN | 0.3 |
| VATB2_HUMAN | 0.13 |
| ANX11_HUMAN | 0.94 |
| ARHG1_HUMAN | 0.9 |
| ERF3A_HUMAN | 1 |
| AIMP2_HUMAN | 1 |
| RAN_HUMAN | 1.11 |
| HNRPD_HUMAN | 1.07 |
| DDX6_HUMAN | 0.72 |
| FUBP1_HUMAN | 1 |
| ANM5_HUMAN | 0 |
| PFD4_HUMAN | 1 |
| MDHM_HUMAN | 0.12 |
| PRPS2_HUMAN | 0.53 |
| TBCA_HUMAN | 0.92 |
| UBP5_HUMAN | 1 |
| FAF1_HUMAN | 1 |
| SPCS_HUMAN | 0.08 |
| SATB1_HUMAN | 0.94 |
| RS11_HUMAN | 0.65 |
| RM37_HUMAN | 1.17 |
| MYO1G_HUMAN | 0.22 |
| DRG1_HUMAN | 0.95 |
| IDHP_HUMAN | 0.08 |
| ADRM1_HUMAN | 0.08 |
| COX41_HUMAN | 1 |
| SF3B2_HUMAN | 0.04 |
| P5CR1_HUMAN | 0.63 |

| | |
|---|---|
| RD23B_HUMAN | 0.42 |
| DNJC9_HUMAN | 1 |
| ECH1_HUMAN | 0.83 |
| SYRC_HUMAN | 0.06 |
| RBMX_HUMAN | 1.11 |
| OLA1_HUMAN | 1 |
| IMDH1_HUMAN | 3.46 |
| RL26_HUMAN | 0.07 |
| RL5_HUMAN | 0.12 |
| PRP8_HUMAN | 1.13 |
| VATG1_HUMAN | 1 |
| IMA1_HUMAN | 0.65 |
| COPZ1_HUMAN | 0.4 |
| TXD17_HUMAN | 1 |
| NUP50_HUMAN | 0.11 |
| RBM4B_HUMAN | |
| SERC_HUMAN | 0.13 |
| ISY1_HUMAN | 0.44 |
| CAPG_HUMAN | 1 |
| GEPH_HUMAN | 0.77 |
| PUR8_HUMAN | 0.82 |
| ARPC2_HUMAN | 0.73 |
| SYCC_HUMAN | 1.07 |
| NAGK_HUMAN | 0.51 |
| DREB_HUMAN | 0.15 |
| ETFA_HUMAN | 0.7 |
| BID_HUMAN | 1 |
| CALX_HUMAN | 1.05 |
| CSN4_HUMAN | 0.17 |
| RS26L_HUMAN | 1 |
| PSMD1_HUMAN | 0.03 |
| ADA_HUMAN | 1.14 |
| ACON_HUMAN | 1 |
| TM109_HUMAN | 0.12 |
| SYMC_HUMAN | 0.89 |
| RL6_HUMAN | 1.04 |
| SYYC_HUMAN | 1.02 |
| RNH2C_HUMAN | 0.55 |
| SYEP_HUMAN | 0.09 |
| BOLA2_HUMAN | 1 |
| ODO2_HUMAN | 0.98 |
| ACTZ_HUMAN | 0.06 |
| AN32A_HUMAN | 1.64 |
| DUT_HUMAN | 0.56 |
| SMD1_HUMAN | 0.74 |
| MARE1_HUMAN | 0.73 |
| CPSF7_HUMAN | 0.82 |
| SPEE_HUMAN | 0.6 |
| BUB3_HUMAN | 1.08 |
| PSME1_HUMAN | 0.5 |
| K2C6A_HUMAN | 0.11 |
| ZN281_HUMAN | 1 |
| K1C14_HUMAN | 0.7 |
| MCA3_HUMAN | 0.16 |
| SRP72_HUMAN | 103.93 |
| TCP4_HUMAN | 0.87 |
| PAIRB_HUMAN | 1.04 |
| GCP3_HUMAN | 3.14 |
| KV109_HUMAN | 1 |
| TAGL2_HUMAN | 6.27 |

| | |
|---|---|
| CSK2B_HUMAN | 8.36 |
| CSK22_HUMAN | 0.98 |
| ATP5L_HUMAN | 9.64 |
| PSB4_HUMAN | 0.15 |
| UBL4A_HUMAN | 0.49 |
| RL10L_HUMAN | 6.18 |
| TBL1R_HUMAN | 0.75 |
| HNRH2_HUMAN | 1 |
| WDR1_HUMAN | 0.96 |
| PP2AB_HUMAN | 1 |
| DCTN2_HUMAN | 0.06 |
| 8ODP_HUMAN | 0.1 |
| PSMD6_HUMAN | 0.07 |
| SUCB2_HUMAN | 1 |
| P5CS_HUMAN | 0.69 |
| PSB1_HUMAN | 0.98 |
| BLM_HUMAN | 1 |
| PUR9_HUMAN | 0.12 |
| DNJA2_HUMAN | 0.72 |
| GCP2_HUMAN | 0.71 |
| MZB1_HUMAN | 1 |
| HYOU1_HUMAN | 1.35 |
| XPO2_HUMAN | 1.14 |
| U2AF2_HUMAN | 0.65 |
| GSTP1_HUMAN | 1.81 |
| UFD1_HUMAN | 0.32 |
| CSN6_HUMAN | 0.03 |
| SYTC_HUMAN | 12.52 |
| RPB3_HUMAN | 0.19 |
| RAB1B_HUMAN | 1 |
| SMC2_HUMAN | 0.05 |
| DNJB1_HUMAN | 0.71 |
| LONM_HUMAN | 0.47 |
| SAE1_HUMAN | 1 |
| U520_HUMAN | 1.1 |
| NSF_HUMAN | 0.11 |
| DGKA_HUMAN | 1.84 |
| ATG3_HUMAN | 1 |
| PLBL2_HUMAN | 1.25 |
| IF6_HUMAN | 1.38 |
| DYHC2_HUMAN | 1 |
| EIF3E_HUMAN | 0.87 |
| U2AF1_HUMAN | 0.17 |
| GIT1_HUMAN | 0.54 |
| K2C71_HUMAN | 0.06 |
| FBRL_HUMAN | 0.19 |
| SF01_HUMAN | 0.84 |
| ZW10_HUMAN | 0.71 |
| ARHG7_HUMAN | 2.47 |
| RFC5_HUMAN | 0 |
| CASP_HUMAN | 1 |
| U5S1_HUMAN | 1.13 |
| EWS_HUMAN | 0.68 |
| DX39B_HUMAN | 1 |
| GNAI3_HUMAN | 0.67 |
| DX39A_HUMAN | 0.16 |
| HAT1_HUMAN | 1 |
| ASNA_HUMAN | 1.01 |
| AP1G1_HUMAN | 1 |
| CDC5L_HUMAN | 1.55 |

| | |
|---|---|
| SF3B1_HUMAN | 1.06 |
| PTBP1_HUMAN | 0.71 |
| SP16H_HUMAN | 0.11 |
| GNAI2_HUMAN | 0.1 |
| RM12_HUMAN | 0.02 |
| PDCD6_HUMAN | 1.47 |
| PPIB_HUMAN | 0.17 |
| IF4B_HUMAN | 1 |
| LARP1_HUMAN | 1 |
| PSMD7_HUMAN | 0.11 |
| ELP5_HUMAN | 0.63 |
| DEOC_HUMAN | 0.43 |
| COTL1_HUMAN | 1.14 |
| RUXGL_HUMAN | 1 |
| K1C15_HUMAN | 9.66 |
| AKAP8_HUMAN | 0.61 |
| MEP50_HUMAN | 0.69 |
| MYH10_HUMAN | 1.43 |
| K2C8_HUMAN | 0.23 |
| SIR2_HUMAN | 0.83 |
| PP1A_HUMAN | 1.05 |
| OSBL7_HUMAN | 1 |
| BZW1_HUMAN | 1 |
| ZAP70_HUMAN | 0.2 |
| KIF11_HUMAN | 0.3 |
| UCHL5_HUMAN | 0.16 |
| FKBP8_HUMAN | 0.36 |
| EI2BG_HUMAN | 1 |
| CCD22_HUMAN | 1 |
| AL9A1_HUMAN | 0.15 |
| PCMD1_HUMAN | 3.18 |
| RINI_HUMAN | 0.95 |
| KLC2_HUMAN | 0.63 |
| LMAN1_HUMAN | 0.9 |
| CSDE1_HUMAN | 0.05 |
| PRS6A_HUMAN | 0.77 |
| SMD2_HUMAN | 1.47 |
| MTDC_HUMAN | 1 |
| EIF3I_HUMAN | 0.15 |
| ECHB_HUMAN | 1.02 |
| ROA0_HUMAN | 0.78 |
| QOR_HUMAN | 0.37 |
| VPS35_HUMAN | 3.52 |
| LV211_HUMAN | 1 |
| SAR1A_HUMAN | 1 |
| PARP1_HUMAN | 0.44 |
| ODO1_HUMAN | 1.09 |
| CECR5_HUMAN | 1 |
| ZYX_HUMAN | 1.23 |
| PABP1_HUMAN | 0.09 |
| CPSF6_HUMAN | 1.14 |
| PSD13_HUMAN | 1.61 |
| TAL1_HUMAN | 0.92 |
| PFKAP_HUMAN | 5.69 |
| RT22_HUMAN | 1 |
| RFC4_HUMAN | 0.15 |
| K1C13_HUMAN | 0.86 |
| FMNL1_HUMAN | 0.12 |
| RT31_HUMAN | 4.31 |
| CSN5_HUMAN | 0.09 |

| | |
|---|---|
| FKBP4_HUMAN | 0.7 |
| SSBP2_HUMAN | 0.67 |
| SYVC_HUMAN | 10.71 |
| OSBL8_HUMAN | 0.98 |
| UBP14_HUMAN | 0.26 |
| RFC3_HUMAN | 0.22 |
| ELMO1_HUMAN | 0.16 |
| IF2A_HUMAN | 1.05 |
| PSA3_HUMAN | 1.89 |
| UBQL1_HUMAN | 1 |
| FNBP1_HUMAN | 1.18 |
| PR40A_HUMAN | 0.85 |
| RBM4_HUMAN | 0.35 |
| DCTN4_HUMAN | 0.24 |
| QCR2_HUMAN | 0.17 |
| PACN2_HUMAN | 0.3 |
| NSF1C_HUMAN | |
| CUL3_HUMAN | 0.95 |
| UBS3A_HUMAN | 0.39 |
| F175B_HUMAN | 0.37 |
| ARK72_HUMAN | 0.32 |
| RCOR3_HUMAN | 0.29 |
| IDE_HUMAN | 0.36 |
| NOLC1_HUMAN | 0.94 |
| GIT2_HUMAN | 0.38 |
| PFKAL_HUMAN | 1.09 |
| CYFP1_HUMAN | 0.43 |
| CYFP2_HUMAN | 1.71 |
| DYR1A_HUMAN | 0.86 |
| CNOT1_HUMAN | 1.29 |
| AP2A2_HUMAN | 1.69 |
| CPZIP_HUMAN | 0.28 |
| SARM1_HUMAN | 0.83 |
| TF2H4_HUMAN | 0.36 |
| P66B_HUMAN | 0.32 |
| VWA8_HUMAN | 0.66 |
| CCAR1_HUMAN | 7.91 |
| EMD_HUMAN | 0.37 |
| RL9_HUMAN | 0.94 |
| SNRPA_HUMAN | 1.39 |
| RU2A_HUMAN | 1.14 |
| PDCL3_HUMAN | 3 |
| PRS10_HUMAN | 0.09 |
| SATB2_HUMAN | 1 |
| HNRPR_HUMAN | 0.12 |
| ACPH_HUMAN | 0.86 |
| ILF2_HUMAN | 0.87 |
| SNF5_HUMAN | 5.94 |
| CDK6_HUMAN | 1.74 |
| CLCA_HUMAN | 0.91 |
| RT29_HUMAN | 1.14 |
| HLAH_HUMAN | 1 |
| AATC_HUMAN | 1 |
| KHDR1_HUMAN | 1.01 |
| CAPZB_HUMAN | 0.58 |
| UBF1_HUMAN | 1.26 |
| COX5A_HUMAN | 1 |
| HYES_HUMAN | 1 |
| AN32E_HUMAN | 1 |
| SC24C_HUMAN | 2.05 |

| | |
|---|---|
| AP3B1_HUMAN | 3.98 |
| ERO1A_HUMAN | 1 |
| LRRF1_HUMAN | 1.23 |
| PFD2_HUMAN | 0.15 |
| DDI2_HUMAN | 0.26 |
| PTPA_HUMAN | 1 |
| NUCB2_HUMAN | 0.24 |
| RM40_HUMAN | 3.33 |
| DC1I2_HUMAN | 1 |
| ILF3_HUMAN | 0.95 |
| SMCA4_HUMAN | 0.23 |
| PSMD8_HUMAN | 0.25 |
| STML2_HUMAN | 1.15 |
| VPS29_HUMAN | 0.2 |
| PSMD4_HUMAN | 0.72 |
| RL13_HUMAN | 0.1 |
| MYLK2_HUMAN | 0.71 |
| DDB1_HUMAN | 0.93 |
| EIF3L_HUMAN | 0.75 |
| TKT_HUMAN | 0.07 |
| EVL_HUMAN | 0.71 |
| ECHA_HUMAN | 0.13 |
| DEFI6_HUMAN | 1 |
| MYH14_HUMAN | 1 |
| DDX23_HUMAN | 1.19 |
| ODPA_HUMAN | 1.31 |
| DDX3X_HUMAN | 0.04 |
| IF4A3_HUMAN | 0.72 |
| RPIA_HUMAN | 1.02 |
| TOP3A_HUMAN | 1 |
| EIF3G_HUMAN | 0.88 |
| POMP_HUMAN | 0.42 |
| VDAC3_HUMAN | 1 |
| PA1B3_HUMAN | 0.09 |
| GDIA_HUMAN | 1 |
| KV202_HUMAN | 1 |
| UB2L3_HUMAN | 1 |
| GMFG_HUMAN | 0.26 |
| GNL1_HUMAN | 1 |
| RPB2_HUMAN | 0.46 |
| AIP_HUMAN | 0.58 |
| SRSF6_HUMAN | 0.07 |
| CAN1_HUMAN | 0.71 |
| AMPD2_HUMAN | 0.75 |
| SEP11_HUMAN | 0.91 |
| UBE2N_HUMAN | 1 |
| SMC4_HUMAN | 1 |
| MAP4_HUMAN | 0.22 |
| LEG1_HUMAN | 1 |
| SC11A_HUMAN | 1 |
| SYFB_HUMAN | 0.17 |
| SSBP3_HUMAN | 1.15 |
| GRB2_HUMAN | 1.7 |
| CARM1_HUMAN | 0.63 |
| AP1M1_HUMAN | 0.8 |
| AGO2_HUMAN | 0.34 |
| TBAL3_HUMAN | 1 |
| TLN1_HUMAN | 1.12 |
| COPB2_HUMAN | 0.8 |
| DLDH_HUMAN | 0.71 |

| | |
|---|---|
| RT23_HUMAN | 2.28 |
| TOP1_HUMAN | 1.23 |
| HNRL1_HUMAN | 0 |
| 1B07_HUMAN | 1 |
| CNN2_HUMAN | 0.91 |
| H1X_HUMAN | 1 |
| ARHG6_HUMAN | 1.49 |
| EIF3K_HUMAN | 0.03 |
| DDRGK_HUMAN | 1 |
| CCAR2_HUMAN | 1.32 |
| HP1B3_HUMAN | 0.63 |
| RPN2_HUMAN | 0.67 |
| ROAA_HUMAN | 0.86 |
| IKZF3_HUMAN | 1 |
| VDAC1_HUMAN | 1.13 |
| AP2B1_HUMAN | 0.97 |
| COPE_HUMAN | 0.78 |
| PP4C_HUMAN | 0.67 |
| PARK7_HUMAN | 0.19 |
| CAB39_HUMAN | 1 |
| TNPO1_HUMAN | 1.33 |
| MTA2_HUMAN | 0.1 |
| PSME2_HUMAN | 0.13 |
| RS5_HUMAN | 0.16 |
| BZW2_HUMAN | 0.19 |
| PDS5A_HUMAN | 0.46 |
| RHG04_HUMAN | 0.13 |
| PGAM1_HUMAN | 1.76 |
| ASNS_HUMAN | 1 |
| CRKL_HUMAN | 1 |
| PUF60_HUMAN | 0.84 |
| SODC_HUMAN | 1 |
| CALR_HUMAN | 0.09 |
| KV106_HUMAN | 1 |
| PSA6_HUMAN | 1.32 |
| NSUN2_HUMAN | 0.76 |
| LMNB2_HUMAN | 1 |
| AATM_HUMAN | 1.2 |
| PTCD3_HUMAN | 0.18 |
| VAT1_HUMAN | 0.21 |
| LRC59_HUMAN | 0.65 |
| KLC1_HUMAN | 0.86 |
| PDIA4_HUMAN | 1 |
| AAKG1_HUMAN | 0.54 |
| PRDX6_HUMAN | 4.37 |
| TBCB_HUMAN | 4.16 |
| NDUA4_HUMAN | 1 |
| SYSC_HUMAN | 1 |
| DC1L1_HUMAN | 0.58 |
| RANB9_HUMAN | 1.17 |
| PA2G4_HUMAN | 1.22 |
| CASP8_HUMAN | 19.54 |
| EI2BA_HUMAN | 1 |
| HNRPQ_HUMAN | 0.97 |
| NHRF1_HUMAN | 1.07 |
| IF16_HUMAN | 0.71 |
| TCEA1_HUMAN | 1 |
| CATA_HUMAN | 0.14 |
| K1C18_HUMAN | 1 |
| KT33B_HUMAN | 7.56 |

| | |
|---|---|
| PYGB_HUMAN | 1.17 |
| EDC4_HUMAN | 0.98 |
| ICLN_HUMAN | 0.56 |
| ACL6A_HUMAN | 0.79 |
| PEBP1_HUMAN | 1 |
| RB11A_HUMAN | 1 |
| ARPC3_HUMAN | 0.18 |
| COPG2_HUMAN | 0.54 |
| CD2AP_HUMAN | 1.17 |
| AP2A1_HUMAN | 0.21 |
| LASP1_HUMAN | 0.11 |
| RIR2_HUMAN | 0.07 |
| NUP62_HUMAN | 0.32 |
| LMNB1_HUMAN | 1.69 |
| SMCE1_HUMAN | 0.78 |
| YTHD2_HUMAN | 1.34 |
| CALU_HUMAN | 1.1 |
| SSRD_HUMAN | 4.02 |
| IQGA1_HUMAN | 1.1 |
| K1C16_HUMAN | 0.05 |
| SKP1_HUMAN | 0.8 |
| TTC28_HUMAN | 1 |
| K1C19_HUMAN | 0.09 |
| CDK9_HUMAN | 0.75 |
| NUP54_HUMAN | 0.63 |
| CAND1_HUMAN | 0.1 |
| PSB6_HUMAN | 1.13 |
| RAB5C_HUMAN | 1 |
| PCH2_HUMAN | 1.16 |
| ADDA_HUMAN | 1.18 |
| RMXL1_HUMAN | 1 |
| DYN2_HUMAN | 1.18 |
| ESYT2_HUMAN | 0.54 |
| IGBP1_HUMAN | 0.18 |
| KCAB2_HUMAN | 0.86 |
| HTF4_HUMAN | 1.49 |
| SND1_HUMAN | 2.29 |
| LPPRC_HUMAN | 0.86 |
| UBP2L_HUMAN | 0.92 |
| CREB1_HUMAN | 1 |
| CNDP2_HUMAN | 0.75 |
| SGTA_HUMAN | 0.41 |
| ODBA_HUMAN | 1.23 |
| LMNA_HUMAN | 1 |
| PDC6I_HUMAN | 0.07 |
| IMA4_HUMAN | 0.58 |
| ATPO_HUMAN | 0.85 |
| HCLS1_HUMAN | 1 |
| BACH_HUMAN | 0.78 |
| COR1C_HUMAN | 0.19 |
| PSB8_HUMAN | 0.24 |
| FUS_HUMAN | 1.05 |
| T2FA_HUMAN | 1 |
| WASF2_HUMAN | 0.31 |
| DPOD2_HUMAN | 0.21 |
| CND2_HUMAN | 0.7 |
| CUL5_HUMAN | 0.48 |
| CUL4A_HUMAN | 0.98 |
| ADAS_HUMAN | 0.43 |
| DSG1_HUMAN | 1 |

| | |
|---|---|
| 2A5D_HUMAN | 0.14 |
| NDUS1_HUMAN | 1 |
| PAF1_HUMAN | 0.43 |
| PRP6_HUMAN | 0.18 |
| RMI1_HUMAN | 1 |
| TADBP_HUMAN | 1 |
| EIF3B_HUMAN | 0.12 |
| HTSF1_HUMAN | 1 |
| IF4G1_HUMAN | 0.07 |
| TRXR1_HUMAN | 1 |
| FLNB_HUMAN | 0.14 |
| 2ABA_HUMAN | 1.23 |
| ATX2L_HUMAN | 0.84 |
| CDC37_HUMAN | 0.96 |
| REVERSE19825 | 1 |
| MIC60_HUMAN | 1.53 |
| CLPX_HUMAN | 0.71 |
| ERF1_HUMAN | 1.12 |
| NUP93_HUMAN | 22.78 |
| DNJA3_HUMAN | 0.97 |
| PCY2_HUMAN | 0.57 |
| IDH3B_HUMAN | 0.36 |
| BLMH_HUMAN | 0.09 |
| KAP1_HUMAN | 0.29 |
| GLSK_HUMAN | 0.66 |
| ARMC8_HUMAN | 0.17 |
| RCOR1_HUMAN | 0.28 |
| PAPS1_HUMAN | 0.26 |
| PP6R1_HUMAN | 0.83 |
| AP1B1_HUMAN | 0.74 |
| EXOS8_HUMAN | 0.23 |
| VASP_HUMAN | 0.2 |
| WDR61_HUMAN | 0.21 |
| RAGP1_HUMAN | 0.87 |
| MYPT1_HUMAN | 0.24 |
| NU107_HUMAN | 0.35 |
| ADHX_HUMAN | 0.29 |
| BCS1_HUMAN | 0.46 |
| WDR82_HUMAN | 0.44 |
| EIF3H_HUMAN | 0.73 |
| EXOC2_HUMAN | 1 |
| RUNX1_HUMAN | 1 |
| CAPR1_HUMAN | 1.03 |
| CCDC6_HUMAN | 0.27 |
| SYWC_HUMAN | 1 |
| PAK2_HUMAN | 0.15 |
| DPOD3_HUMAN | 0.33 |
| GRWD1_HUMAN | 0.2 |
| MARE2_HUMAN | 0.43 |
| SC23A_HUMAN | 0.44 |
| IF2B3_HUMAN | 0.53 |
| EHD1_HUMAN | 0.33 |
| BAG5_HUMAN | 0.27 |
| DCUP_HUMAN | 0.69 |
| CDV3_HUMAN | 0.1 |
| TBK1_HUMAN | 0.4 |
| DDX46_HUMAN | 0.41 |
| PACS1_HUMAN | 1.55 |
| STT3A_HUMAN | 1.62 |
| DNLI1_HUMAN | 1.4 |

| | |
|---|---|
| MBB1A_HUMAN | 0.26 |
| THMS1_HUMAN | 0.79 |
| AN32B_HUMAN | 2.18 |
| REQU_HUMAN | 0.27 |
| ANXA2_HUMAN | 1 |
| OXSR1_HUMAN | 0.7 |
| DNMT1_HUMAN | 1.65 |
| BAG2_HUMAN | 1.06 |
| AP3S1_HUMAN | 0.27 |
| MK01_HUMAN | 0.49 |
| FEN1_HUMAN | 1.42 |
| RAB14_HUMAN | 1 |
| DHE3_HUMAN | 0.8 |
| IDHC_HUMAN | 0.49 |
| PSMD5_HUMAN | 0.81 |
| DPP3_HUMAN | 1 |
| ARP2_HUMAN | 0.1 |
| PP1B_HUMAN | 2.25 |
| RM50_HUMAN | 0.89 |
| SRP54_HUMAN | 0.1 |
| PDIP2_HUMAN | 0.34 |
| MAOM_HUMAN | 0.7 |
| PPM1G_HUMAN | 0.14 |
| SPAT5_HUMAN | 0 |
| ODB2_HUMAN | 0.03 |
| TYSY_HUMAN | 0.38 |
| PPP6_HUMAN | 0.65 |
| PAP1M_HUMAN | 1 |
| NP1L4_HUMAN | 0.84 |
| EXOS9_HUMAN | 0.16 |
| RT34_HUMAN | 0.46 |
| K1C25_HUMAN | 0.63 |
| EIF3C_HUMAN | 0.12 |
| EIF3A_HUMAN | 1.31 |
| NELFB_HUMAN | 0.25 |
| RS4Y2_HUMAN | 1 |
| CSK_HUMAN | 0.64 |
| SRSF4_HUMAN | 1 |
| STIM1_HUMAN | 0.43 |
| SYAC_HUMAN | 0.47 |
| VATA_HUMAN | 0.26 |
| TTF2_HUMAN | 1 |
| SNAA_HUMAN | 0.51 |
| SFXN1_HUMAN | 1 |
| ERP29_HUMAN | 1.17 |
| OAT_HUMAN | 0.69 |
| GLRX3_HUMAN | 0.32 |
| DDX41_HUMAN | 0.44 |
| SR140_HUMAN | 1.28 |
| AKP8L_HUMAN | 0.69 |
| AAAT_HUMAN | 0.3 |
| 2AAB_HUMAN | 0.63 |
| DNJC7_HUMAN | 0.17 |
| CTBP1_HUMAN | 1.34 |
| DHB4_HUMAN | 1.02 |
| RENT1_HUMAN | 1.27 |
| ARI5B_HUMAN | 1 |
| K1C12_HUMAN | 1 |
| PFD6_HUMAN | 1 |
| TRIPB_HUMAN | 1 |

| | |
|---|---|
| SRP68_HUMAN | 0.11 |
| VDAC2_HUMAN | 1.08 |
| TMOD3_HUMAN | 0.19 |
| HNRDL_HUMAN | 0.43 |
| VP26B_HUMAN | 0.25 |
| RASL3_HUMAN | 0.56 |
| KLHL7_HUMAN | 0.26 |
| PMS2_HUMAN | 1 |
| HNRPL_HUMAN | 7.54 |
| SCYL2_HUMAN | 0.83 |
| KPRA_HUMAN | 1 |
| GLU2B_HUMAN | 0.1 |
| PYRG1_HUMAN | 0.74 |
| SGT1_HUMAN | 0.13 |
| PUR2_HUMAN | 0.2 |
| EXOSX_HUMAN | 0.28 |
| SRSF3_HUMAN | 0.98 |
| DDX21_HUMAN | 42.29 |
| STK4_HUMAN | 2.88 |
| ELP3_HUMAN | 0.59 |
| PP1G_HUMAN | 1.1 |
| UB2V1_HUMAN | 0.06 |
| BAG6_HUMAN | 0.28 |
| AIFM1_HUMAN | 0.04 |
| DBNL_HUMAN | 1 |
| PRDX5_HUMAN | 0.14 |
| DCTN1_HUMAN | 0.11 |
| SMRC2_HUMAN | 1.5 |
| REVERSE12748 | 1 |
| TOIP1_HUMAN | 1 |
| XPOT_HUMAN | 1 |
| SC31A_HUMAN | 0.46 |
| LPXN_HUMAN | 0.85 |
| TLE3_HUMAN | 0.38 |
| UBAP2_HUMAN | 1 |
| PSA_HUMAN | 1 |
| LEMD2_HUMAN | 1 |
| G3BP1_HUMAN | 0.65 |
| MLH1_HUMAN | 0.06 |
| NASP_HUMAN | 0.19 |
| LEF1_HUMAN | 0.26 |
| 1B41_HUMAN | 2.97 |
| WRIP1_HUMAN | 0.79 |
| HGH1_HUMAN | 0.83 |
| RFC1_HUMAN | 0.39 |
| ROCK2_HUMAN | 1.1 |
| HPBP1_HUMAN | 0.31 |
| PI42A_HUMAN | 0.42 |
| AHSA1_HUMAN | 0.06 |
| ARF4_HUMAN | 1 |
| ABCE1_HUMAN | 0.3 |
| XPO7_HUMAN | 0.45 |
| LIS1_HUMAN | 1.69 |
| TOP2A_HUMAN | 0.71 |
| AP3M1_HUMAN | 2.32 |
| DJC13_HUMAN | 0.18 |
| SF3A2_HUMAN | 1 |
| ZCCHV_HUMAN | 0.74 |
| URP2_HUMAN | 0.19 |
| UB2V2_HUMAN | 0.17 |

| | |
|---|---|
| TIA1_HUMAN | 1.45 |
| OSGEP_HUMAN | 0.4 |
| IKZF1_HUMAN | 3.25 |
| SNUT1_HUMAN | 0.39 |
| TWF2_HUMAN | 0.23 |
| NDUS2_HUMAN | 6.77 |
| UBP11_HUMAN | 0.96 |
| SART3_HUMAN | 1 |
| RCC2_HUMAN | 0.28 |
| PSMG1_HUMAN | 2.09 |
| KDM1A_HUMAN | 1.58 |
| SYF1_HUMAN | 0.95 |
| EI2BE_HUMAN | 2.46 |
| RCN1_HUMAN | 5 |
| AP2M1_HUMAN | 1.03 |
| AGO1_HUMAN | 0.9 |
| ESYT1_HUMAN | 0.17 |
| STRAP_HUMAN | 0.12 |
| SYFA_HUMAN | 0.67 |
| TIM50_HUMAN | 1.06 |
| WASP_HUMAN | 1 |
| C1TM_HUMAN | 0.36 |
| HAX1_HUMAN | 1.4 |
| RT09_HUMAN | 0.18 |
| LBR_HUMAN | 1 |
| CSN2_HUMAN | 0.17 |
| DHX9_HUMAN | 0.1 |
| K1H1_HUMAN | 0.57 |
| K1H2_HUMAN | 0.07 |
| SRRT_HUMAN | 1.07 |
| ARC1B_HUMAN | 0.27 |
| RAB7A_HUMAN | 0.19 |
| BYST_HUMAN | 0.02 |
| LCK_HUMAN | 3.75 |
| EIF2A_HUMAN | 1.06 |
| XRN2_HUMAN | 0.36 |
| AT1B3_HUMAN | 1 |
| IPO5_HUMAN | 8.76 |
| DOCK2_HUMAN | 1.09 |
| RFA1_HUMAN | 0.12 |
| THIL_HUMAN | 0.91 |
| FLII_HUMAN | 1.2 |
| ELP4_HUMAN | 0.98 |
| HS74L_HUMAN | 0.29 |
| SYNC_HUMAN | 0.55 |
| ANXA5_HUMAN | 1 |
| CND1_HUMAN | 1.36 |
| PURA2_HUMAN | 0.75 |
| EXOC4_HUMAN | 0.02 |
| NACC1_HUMAN | 2.19 |
| ELP1_HUMAN | 0.3 |
| CDC73_HUMAN | 0.48 |
| SUCA_HUMAN | 0.35 |
| SK2L2_HUMAN | 0.66 |
| I2BP2_HUMAN | 1 |
| SMC1A_HUMAN | 1 |
| KCD12_HUMAN | 0.65 |
| BRE1B_HUMAN | 1.25 |
| ALDOC_HUMAN | 1 |
| CKAP5_HUMAN | 0.15 |

| | |
|---|---|
| HMHA1_HUMAN | 1.69 |
| PABP2_HUMAN | 1 |
| HMGB2_HUMAN | 1 |
| ELOC_HUMAN | 7.4 |
| DJC10_HUMAN | 0.36 |
| TPP2_HUMAN | 0.27 |
| HNRH3_HUMAN | 1.35 |
| RFC2_HUMAN | 1.17 |
| LCP2_HUMAN | 1 |
| TNR6_HUMAN | 3.99 |
| IF4G2_HUMAN | 0.94 |
| TXND5_HUMAN | 0.4 |
| BCR_HUMAN | 0.47 |
| KIF4A_HUMAN | 1 |
| SH3K1_HUMAN | 1 |
| TCOF_HUMAN | 0.13 |
| RM11_HUMAN | 0.44 |
| SRSF7_HUMAN | 0.88 |
| LS14B_HUMAN | 0.37 |
| PABP4_HUMAN | 1 |
| KIF2C_HUMAN | 0.94 |
| CCNK_HUMAN | 0.54 |
| DCTN3_HUMAN | 0.31 |
| RBM14_HUMAN | 0.44 |
| LETM1_HUMAN | 1 |
| DDX42_HUMAN | 5.53 |
| CUL4B_HUMAN | 0.84 |
| DHX30_HUMAN | 0.37 |
| UGPA_HUMAN | 1 |
| RABL6_HUMAN | 0.94 |
| UGGG1_HUMAN | 0.76 |
| SNX1_HUMAN | 0.33 |
| HCDH_HUMAN | 0.34 |
| EI2BB_HUMAN | 0.34 |
| RBM25_HUMAN | 0.37 |
| PLPL8_HUMAN | 64.68 |
| RFA2_HUMAN | 6.73 |
| ARHG2_HUMAN | 0.26 |
| NOP2_HUMAN | 0.43 |
| K2C79_HUMAN | 3.13 |
| XPO1_HUMAN | 1.06 |
| APT_HUMAN | 0.03 |
| PLCG1_HUMAN | 0.2 |
| EIF3D_HUMAN | 0.18 |
| PYR1_HUMAN | 1 |
| WDR26_HUMAN | 0.27 |
| FA49B_HUMAN | 1 |
| FIP1_HUMAN | 1 |
| SYHC_HUMAN | 0.27 |
| TCRG1_HUMAN | 0.37 |
| DNM1L_HUMAN | 0.57 |
| PUR4_HUMAN | 0.51 |
| DMAP1_HUMAN | 0.57 |
| SCFD1_HUMAN | 1 |
| SDHA_HUMAN | 0.17 |
| SPTN1_HUMAN | 0.17 |
| KV119_HUMAN | 1 |
| CLIC1_HUMAN | 0.22 |
| IPYR2_HUMAN | 1.59 |
| CND3_HUMAN | 3.86 |

| | |
|---|---|
| PO210_HUMAN | 0.06 |
| PAK1_HUMAN | 1 |
| EXOC3_HUMAN | 2.24 |
| PNPT1_HUMAN | 1.75 |
| TTL12_HUMAN | 0.24 |
| NUDT5_HUMAN | 5.34 |
| PRPS1_HUMAN | 1 |
| SC23B_HUMAN | 1.27 |
| RAP1A_HUMAN | 0.52 |
| EP15R_HUMAN | 0.66 |
| RSMB_HUMAN | 1.15 |
| DIM1_HUMAN | 0.45 |
| HAUS5_HUMAN | 0.6 |
| DPOD1_HUMAN | 0.23 |
| USO1_HUMAN | 0.9 |
| KCRU_HUMAN | 1.22 |
| VPS16_HUMAN | 2.24 |
| ROCK1_HUMAN | 0.54 |
| VP33B_HUMAN | 0.22 |
| PP6R3_HUMAN | 0.96 |
| TSSC4_HUMAN | 2.4 |
| CUL2_HUMAN | 1.51 |
| PDIA1_HUMAN | 0.22 |
| CSTF2_HUMAN | 0.3 |
| SPF45_HUMAN | 0.2 |
| ATE1_HUMAN | 0.11 |
| WDHD1_HUMAN | 0.08 |
| PRI2_HUMAN | 0.45 |
| RED_HUMAN | 0.36 |
| IKZF2_HUMAN | 1 |
| NAMPT_HUMAN | 1.64 |
| OSBP1_HUMAN | 1.33 |
| RPAP3_HUMAN | 0.68 |
| RANG_HUMAN | 1 |
| SNUT2_HUMAN | 0.32 |
| RPAB3_HUMAN | 1 |
| SMRC1_HUMAN | 0.38 |
| CLU_HUMAN | 0.55 |
| AQR_HUMAN | 0.37 |
| TOP2B_HUMAN | 0.79 |
| STXB2_HUMAN | 0.7 |
| IPO4_HUMAN | 0.4 |
| IF2P_HUMAN | 1.06 |
| RAVR1_HUMAN | 3.92 |
| MPPB_HUMAN | 1 |
| SNX5_HUMAN | 0.71 |
| SNX2_HUMAN | 0.37 |
| XPP1_HUMAN | 0.33 |
| VAPA_HUMAN | 11.51 |
| MCES_HUMAN | 7.45 |
| MDN1_HUMAN | 1 |
| EI2BD_HUMAN | 0.19 |
| MTMR5_HUMAN | 1.02 |
| ACACA_HUMAN | 1.6 |
| CTR9_HUMAN | 1.96 |
| HCFC1_HUMAN | 0.28 |
| KAPCA_HUMAN | 1.16 |
| FAS_HUMAN | 0.05 |
| WASH6_HUMAN | 0.95 |
| LIPA1_HUMAN | 0.52 |

| | |
|---|---|
| GUAA_HUMAN | 0.37 |
| GLE1_HUMAN | 0.02 |
| UBP7_HUMAN | 0.08 |
| ANXA7_HUMAN | 0.23 |
| SC16A_HUMAN | 5.06 |
| SMC3_HUMAN | 1 |
| CAR11_HUMAN | 1 |
| RM39_HUMAN | 0.31 |
| ZCHC8_HUMAN | 0.81 |
| TRI25_HUMAN | 0.31 |
| RHG15_HUMAN | 0.33 |
| FPPS_HUMAN | 0.27 |
| NAT10_HUMAN | 0.5 |
| THIK_HUMAN | 1 |
| AT1A1_HUMAN | 0.3 |
| REVERSE16327 | 1 |
| APEX1_HUMAN | 1 |
| H3C_HUMAN | 1 |
| BIEA_HUMAN | 0.56 |
| CSN3_HUMAN | 5.73 |
| MATR3_HUMAN | 0.98 |
| CPSF3_HUMAN | 1 |
| GAPD1_HUMAN | 0.65 |
| FA21C_HUMAN | 1.15 |
| UBR5_HUMAN | 1.74 |

**Supplementary Table 1:** List of FADD-TAP interacting proteins identified by label-free LC-MS/MS; fold-change in amount of proteins following anti-CD95 treatment. Source data for Volcano plot shown in Figure 2A.

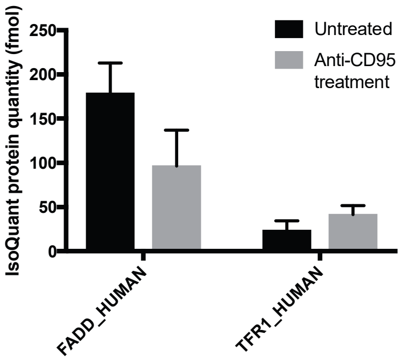


**Supplementary Figure 1** - Label-free quantitation of amount of FADD and Transferrin Receptor 1 (TfR1) detected in FADD-TAP cells +/- anti-CD95 treatment by mass spectrometry (n=3 biological repeats, error bars = SEM).

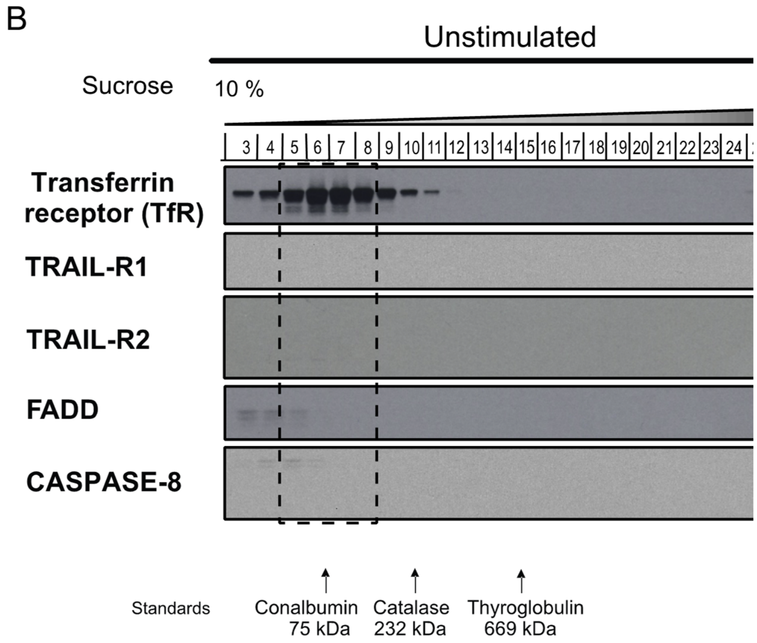


**Supplementary Figure 2** – Western blot analysis of unstimulated BJAB cell lysates separated by buoyant density on 10-45% sucrose density gradient. Note, TfR1 elutes at it's native molecular weight of ~80 kDa (Fractions 5-8). Molecular weight standards were run on a separate gradient and the positions shown are representative of three independent experiments.

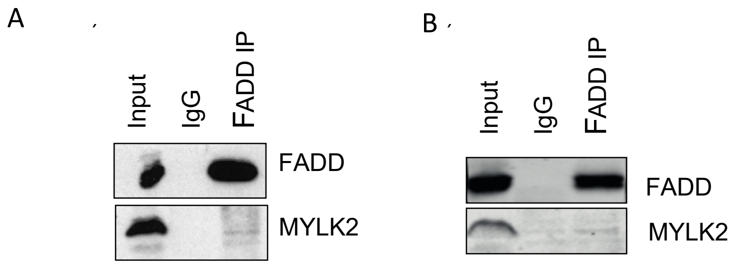


**Supplementary Figure 3** – Analysis of immunoprecipiation of FADD from A) BJAB and B) Jurkat cells, western blotting for FADD and MYLK2.

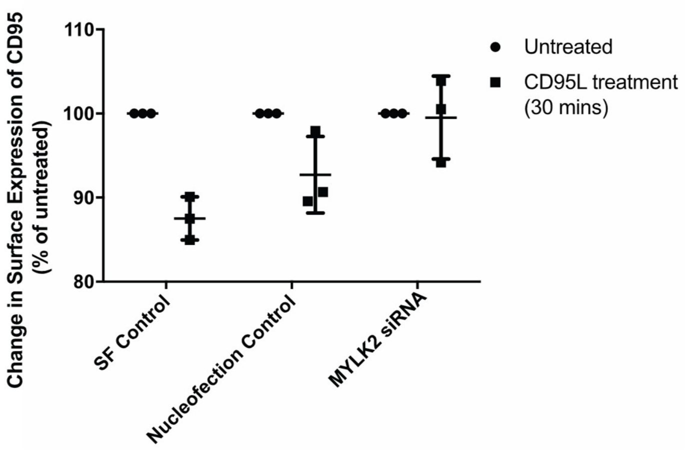


**Supplementary Figure 4** – FACS analysis of surface expression of CD95 following 30 minutes CD95L treatment. SF control = untransfected control, NF = nucleofection control, MYLK2 siRNA = 24hrs MYLK2 knockdown prior to treatment. Surface expression expressed as a percentage compared to the untreated control for each condition. (n=3, Mean +/- SEM).
